# Fission yeast Swi2 designates cell-type specific donor and stimulates Rad51-driven strand exchange

**DOI:** 10.1101/2022.11.22.517464

**Authors:** Takahisa Maki, Geneviève Thon, Hiroshi Iwasaki

## Abstract

A haploid of the fission yeast *Schizosaccharomyces pombe* expresses either the P or M mating-type, determined by the active, euchromatic, *mat1* cassette. Mating-type is switched by Rad51-driven gene conversion of *mat1* using a heterochromatic donor cassette, *mat2-P* or *mat3-M*. The Swi2-Swi5 complex, a mating-type switching factor, is central to this process by designating a preferred donor in a cell-type-specific manner. Swi2-Swi5 selectively enables one of two *cis*acting recombination enhancers, *SRE2* adjacent to *mat2-P* or *SRE3* adjacent to *mat3-M*. Here, we identified two functionally important motifs in Swi2, a Swi6 (HP1 homolog)-binding site and two DNA-binding AT-hooks. Genetic analysis demonstrated that the AT-hooks were required for Swi2 localization at *SRE3* to select the *mat3-M* donor in P cells, while the Swi6-binding site was required for Swi2 localization at *SRE2* to select *mat2-P* in M cells. In addition, the Swi2-Swi5 complex promoted Rad51-driven strand exchange *in vitro*. Taken together, our results show how the Swi2-Swi5 complex would localize to recombination enhancers through a cell-type specific binding mechanism and stimulate Rad51-driven gene conversion at the localization site.

## Introduction

Homothallic strains of the fission yeast *Schizosaccharomyces pombe* (*h^90^*) are either of the P or M mating-type and switch their mating type during vegetative growth (mating-type switching: MTS). The mating type is determined by the transcriptionally active *mat1* cassette at the *mat* locus and is switched by a gene conversion event of the *mat1* cassette using a donor in the heterochromatic silent region, *mat2-P* or *mat3-M*. P cells (*mat1-P*) and M cells (*mat1-M*) preferentially utilize *mat3-M* and *mat2-P* as a donor for gene conversion, respectively (Klar *et al*, 2014).

The chromosome organization of the *mat* locus is shown in Figure 1. The *mat1* cassette expresses either P or M information, *Pi* and *Pc* genes in P cells (*mat1-P*) or *Mi* and *Mc* genes in M cells (*mat1-M*), which determines the mating type of a haploid cell (Kelly *et al*, 1988). The *mat2* and *mat3* cassettes are located in the 20 kb region between *IR-L* and *IR-R*, which is ~15 kb from *mat1*. The *mat2* and *mat3* cassettes contain P (*mat2-P*) and M (*mat3-M*) information, respectively, and are silenced by heterochromatin. Each *mat* cassette is flanked by short homology boxes, *H1* and *H2*, which are used for Rad51-mediated homology-driven gene conversion during MTS (Klar *et al*., 2014). Two *cis*-acting <u>S</u>wi2-dependent <u>r</u>ecombination <u>e</u>nhancers, *SRE2* and *SRE3*, are located adjacent to the *H1* box of *mat2-P* and *mat3-M*, respectively. These two *SREs* play an important role in the donor choice step in each cell type during MTS (Jakociunas *et al*, 2013; Jia *et al*, 2004; Yu *et al*, 2012).

**Figure 1.**
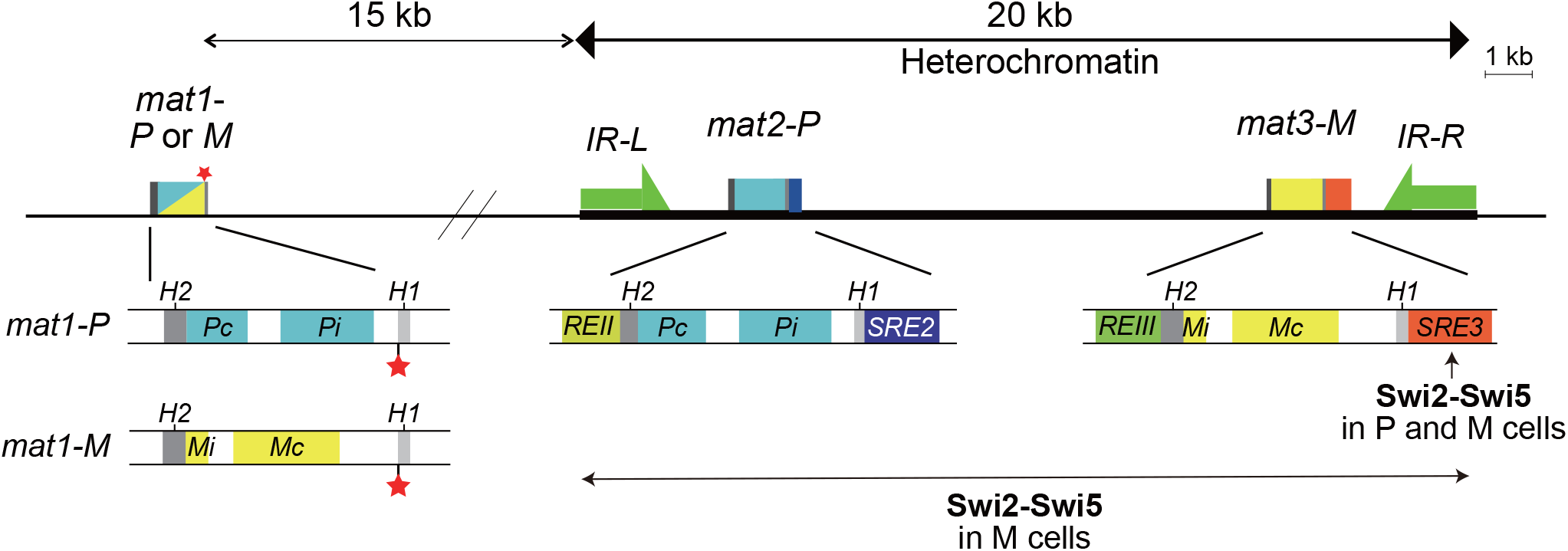
MTS in *S. pombe*. Simple schematic representation of the mating-type region in the *h^90^* strain. The switchable *mat1* cassette contains P (blue; *Pc* and *Pi*) or M (yellow; *Mi* and *Mc*) mating-type genes. The *mat2-P* and *mat3-M* donor cassettes are located in a heterochromatin region between *IR-L* and *IR-R* repeats and their expression is repressed. *REII* and *REIII* stand for Repressor Element II and III, respectively. The *SRE2* and *SRE3* elements are located near the *mat2-P* and *mat3-M* cassettes, respectively. Each cassette is flanked by the *H1* and *H2* homology boxes. The imprinting site at the *mat1-H1* junction is indicated by a red asterisk. Imprinting is converted to a DSB. The MTS factor Swi2-Swi5 localizes to *SRE3* in P cells, but localizes over the *mat* heterochromatin locus in M cells.

MTS is initiated by the introduction of an imprint at the junction of *mat1* and the *H1* box during lagging strand synthesis in DNA replication (Dalgaard & Klar, 1999, 2000). This imprint has been suggested to be a nick or incorporation of two ribonucleotides (Kaykov & Arcangioli, 2004; Singh *et al*, 2019; Singh, 2019; Vengrova & Dalgaard, 2006), and a daughter cell that inherits the imprint is converted to a ‘switchable’ state. On the other hand, a daughter cell that inherits the *de novo* leading strand is unable to switch cell type and is referred to as “unswitchable” (Arcangioli, 1998; Dalgaard & Klar, 1999; Miyata & Miyata, 1981). During leading strand synthesis in a granddaughter cell, the imprint is converted to a single-ended DNA double-strand break (DSB), which triggers the gene conversion event driven by the homologous recombination (HR) protein Rad51 (Roseaulin *et al*, 2008; Yamada-Inagawa *et al*, 2007).

The following model can be put forward to explain gene conversion for MTS driven by Rad51. The DSB end is resected to produce single-stranded DNA (ssDNA) of the *H1* box in the *mat1* cassette, onto which the Rad51 protein forms a nucleoprotein filament (Roseaulin *et al*., 2008). Subsequently, the resultant Rad51 filament searches the *H1* box of the *mat2-P* or *mat3-M* cassette, and then the ssDNA invades either the *H1* box of *mat2-P* or *mat3-M*. This process does not occur randomly. There are highly regulated donor choice mechanisms involving *SRE2* and *SRE3* (Jakociunas *et al*., 2013; Jia *et al*., 2004; Yu *et al*., 2012). The *SRE2* element is required to select *mat2-P* as a donor in M cells, while the *SRE3* element is required to select *mat3-M* as a donor in P cells (Figure 1) (Jakociunas *et al*., 2013). After strand invasion, the DSB is repaired by a synthesis-dependent strand annealing mechanism using the *H2* box (Yamada-Inagawa *et al*., 2007).

Ten MTS genes named *swi1–swi10* were initially identified (Egel *et al*, 1984). Among them, *swi2^+^, swi5^+^* and *swi6^+^* are essential for the normal donor choice mechanism (Thon *et al*, 2019). Swi6 is a homolog of heterochromatin protein 1 (HP1) in other organisms and localizes to heterochromatic regions such as centromeres, telomeres, and the *mat* locus by recognizing histone H3K9 methylation (Allshire & Ekwall, 2015). At the *mat* locus, Swi6 is distributed between the inverted repeats *IR-L* and *IR-R* in both mating types (Noma *et al*, 2001; Thon *et al*, 2002). Interestingly, in the absence of Swi6, *h^90^* cells prefer to use the *mat3-M* donor over the *mat2-P* donor (Jakociunas *et al*., 2013; Jia *et al*., 2004; Thon & Klar, 1993).

Yeast two-hybrid assays have suggested that Swi2 and Swi5 form a complex and that Swi2 also binds to Swi6 (Akamatsu *et al*, 2003). In P cells, the Swi2-Swi5 complex localizes only to *SRE3*, independently of Swi6, which is believed to contribute to the *mat3-M* donor choice. On the other hand, in M cells, the Swi2-Swi5 complex not only localizes to *SRE3*, but also to the heterochromatic region between *IR-L* and *IR-R* dependent on Swi6 (Jia *et al*., 2004). The Swi2 and Swi5 protein levels are increased in M cells due to stimulation of *swi2* and *swi5* gene transcription by the M cell-specific factor mat1-Mc and the CENP-B homolog Abp1 (Matsuda *et al*, 2011). The localization mechanism of the Swi2-Swi5 complex in M cells has been suggested to start from *SRE3* via the physical interaction between Swi2 and Swi6 (Jia *et al*., 2004), although Swi2 has a function at *SRE2* regardless of *SRE3* to facilitate the *mat2-P* donor choice (Jakociunas *et al*., 2013). Swi2 interacts with Rad51 (Akamatsu *et al*., 2003) and therefore it has been hypothesized that the Swi2-Swi5 complex recruits the Rad51 filament formed on the *H1* box of the *mat1* cassette to *SRE2* or *SRE3* in a cell-type specific manner (Thon *et al*., 2019). In addition to heterochromatin formation and the Swi2-Swi5 complex, we recently found that the euchromatic factors Set1C and HULC also affect MTS (Maki *et al*, 2018). Based on genetic analysis, we proposed that Set1C and HULC in M cells inhibit the *mat3-M* donor choice at *SRE3* (Esquivel-Chavez *et al*, 2022).

Swi5 forms a stable complex with another partner, Sfr1. Cells deficient for *sfr1* are defective in recombination repair but exhibit normal MTS. On the other hand, *swi2* mutants have normal DNA repair activity. *swi5*-deficient mutants show defects in both MTS and DNA repair. The Swi5-Sfr1 complex stimulates Rad51-driven strand exchange activity *in vitro* by acting as a recombination activator (Akamatsu *et al*, 2007; Haruta *et al*, 2006; Kurokawa *et al*, 2008). The C-terminal part of Swi2 shares sequence homology with the C-terminal part of Sfr1, which forms an interface with Rad51; therefore, the Swi2-Swi5 complex is also expected to activate Rad51.

In this study, we identified two functionally important regions, the Swi6-binding site and two tandem AT-hook motifs, in Swi2. Swi6-binding site and AT-hook mutants had an MTS-deficient phenotype: Swi6-binding site mutants were biased toward M cells, while AT-hook mutants were biased toward P cells. This result indicates that the interaction between Swi2 and Swi6 is important for the *mat2-P* donor choice in M cells, consistent with a previous proposal (Thon *et al*., 2019), while the AT-hook motifs play a different role in MTS. In addition, we demonstrated that the purified Swi2-Swi5 complex stimulates Rad51-driven strand invasion.

## Results

### Amino acids of Swi2 that are important for interaction with Swi6

Previously, we roughly mapped a Swi6-binding interface to the 210–365 aa region of Swi2 by performing two-hybrid assays in *Saccharomyces cerevisiae* (Akamatsu *et al*., 2003). Here, to identify amino acids in Swi2 that are critically important for Swi6 binding, we first generated a series of C-terminal truncations of Swi2 to more precisely pinpoint the Swi6-binding region. The Swi2 (1–280) truncation mutant, but not the Swi2 (1–260) truncation mutant, interacted with full-length Swi6 in yeast two-hybrid assay, suggesting that 261–280 aa of Swi2 play an essential role in Swi6 binding (Figure 2A). The 270–279 aa are highly conserved among the three fission yeast species *S. pombe, S. octosporus*, and *S. cryophilus* (Figure 2B and Supplementary Figure S1). We next generated a series of plasmids expressing full-length Swi2 in which each of the 270–279 aa, except for A273, was individually replaced by alanine (Figure 2C). Unlike full-length Swi2, *S. cerevisiae* cells expressing Swi2-V270A, -F271A, -V274A, -V275A, -I276A, or -P277A did not grow on 4DO dropout plates indicating poor reporter gene activation and thus defective interaction between the Swi2 mutant proteins and Swi6 (Figure 2D). The Swi2-V274A mutant did not support growth even on 3DO plates where weak interactions can support growth. We conclude that V270, F271, and V274–P277 are essential for Swi6 binding. The Swi2-5A mutant, in which F271, D272, V274, V275, and I276 were all replaced by alanine, exhibited a particularly severe defect in Swi6 binding (Figure 2D).

**Figure 2.**
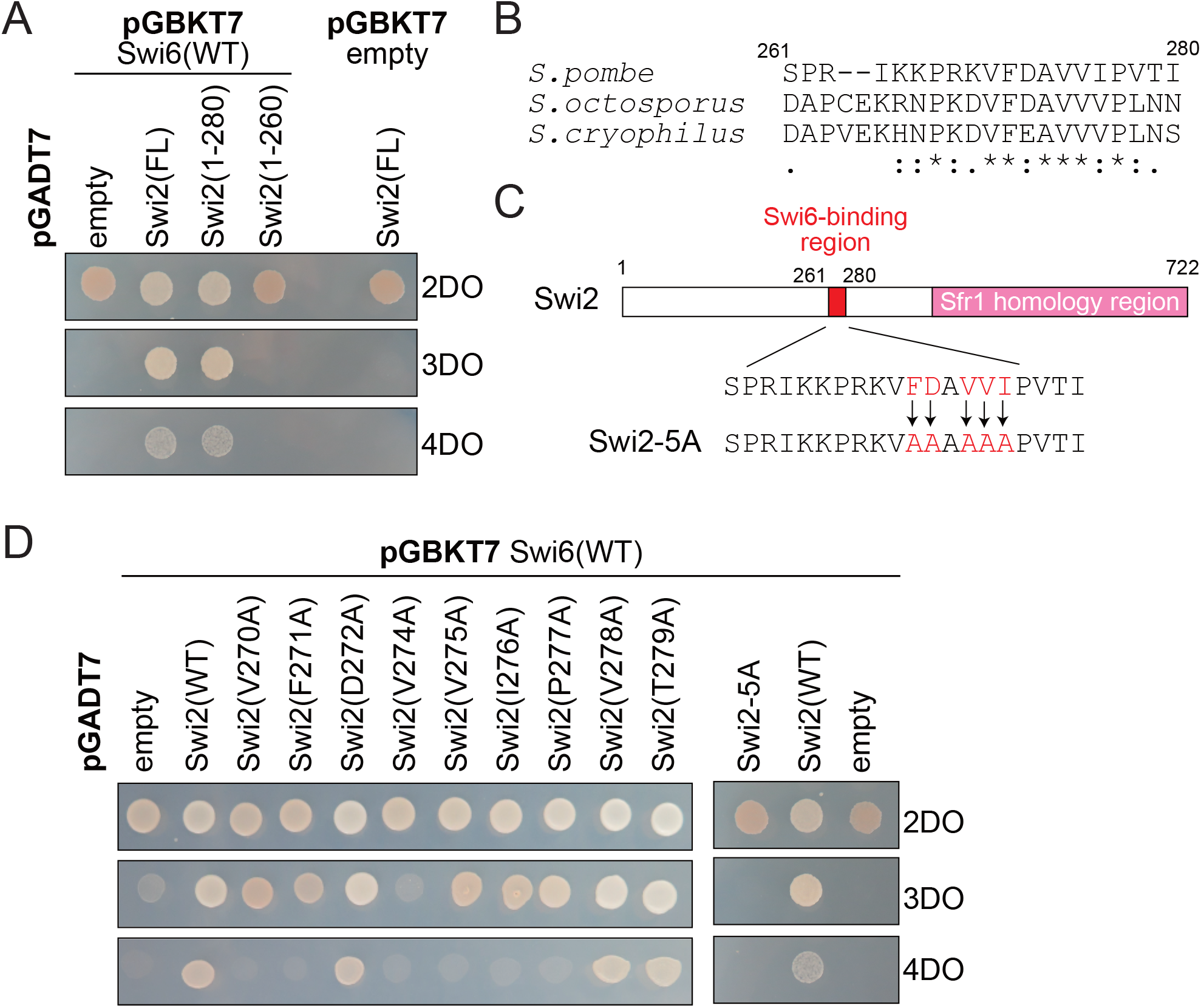
The Swi6-binding site in Swi2. (A) Truncation analysis of Swi2 to map its Swi6-binding region. The yeast two-hybrid assay examined Swi6 interactions with the indicated Swi2 regions (full-length: FL, 1–280 aa, and 1–260 aa). Two-hybrid interactions between Swi2 derivatives and Swi6 were analyzed by spot tests on three types of dropout (DO) plates: control 2DO (SD-leu and -trp) testing for cotransformation by the bait- and prey-encoding plasmids, 3DO (SD-his, -leu, and -trp) testing for activation of the *HIS3* reporter gene by protein interaction, and 4DO (SD-ade, -his, -leu, and - trp) testing for activation of both *HIS3* and *ADE2* reporters by protein interaction. (B) Sequence alignment of the Swi6-binding region in *S. pombe, S. octosporus*, and *S. cryophilus*. (C) The indicated residues of Swi2 in *S. pombe* were mutated to alanine to generate Swi2-5A. (D) The indicated residues of Swi2 were mutated to alanine. The interactions of the Swi2 mutants with Swi6 were examined by a yeast two-hybrid assay as in (A).

### Swi2 has two DNA-binding AT-hook motifs

Sequence analysis revealed that Swi2 proteins from three *Schizosaccharomyces* yeast species share two highly conserved AT-hook motifs located close to the Swi6-binding site (Figure 3A and Supplementary Figure S1). The AT-hook motif is a DNA-binding motif consisting of approximately 15 amino acids with a central GRP consensus sequence rich in positively charged arginine/lysine residues on either side of the core GRP (Aravind & Landsman, 1998; Reeves, 2001). Its core sequence binds to the minor groove of AT-rich DNA (Reeves, 2001).

**Figure 3.**
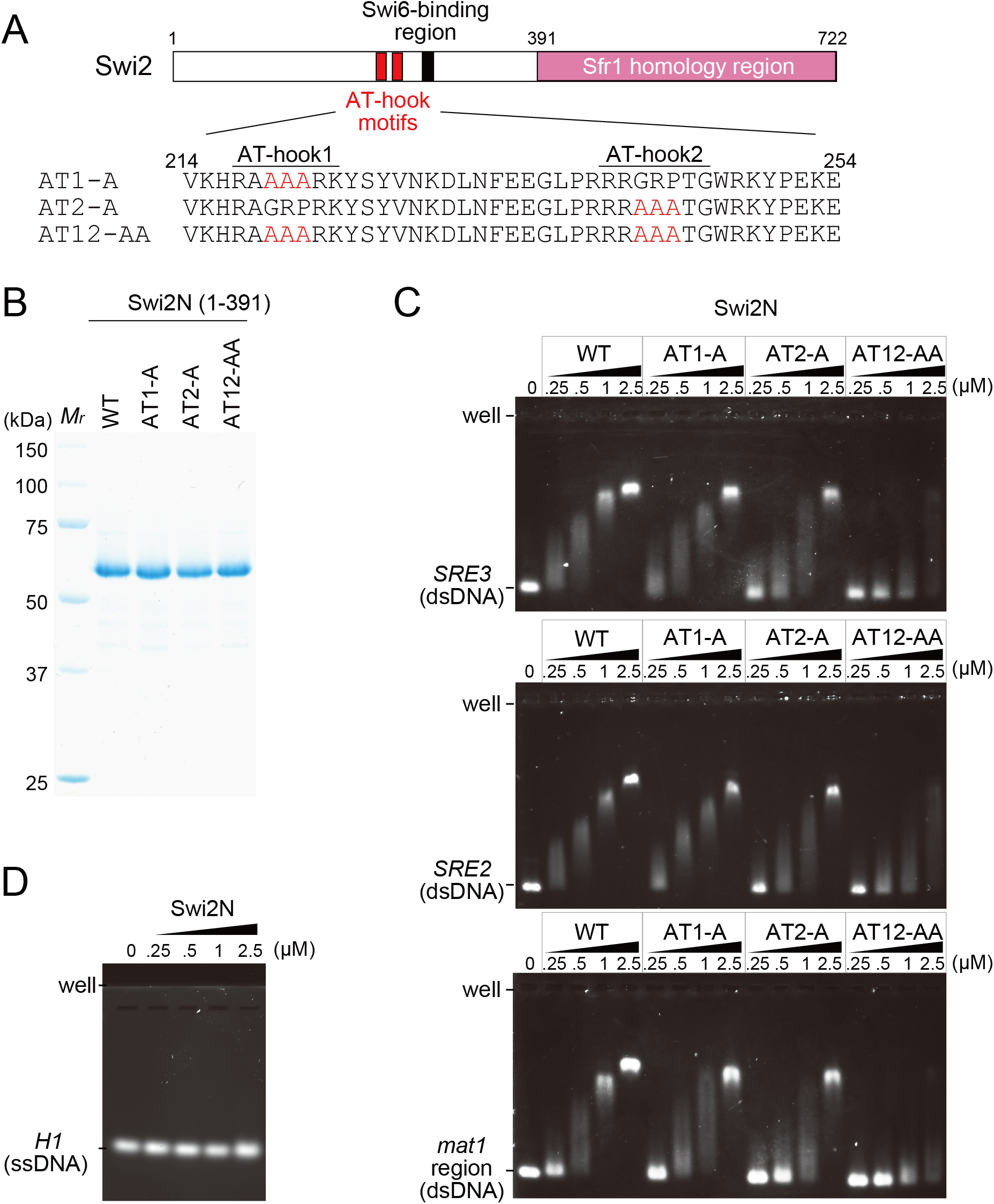
The two AT-hook motifs in Swi2. (A) Schematics of the two AT-hook motifs in Swi2. The consensus sequence, GRP tripeptides, in the two AT-hook motifs were substituted with alanine to generate AT1-A, AT2-A, and AT12-AA. (B) An SDS-PAGE gel of the purified Swi2N derivative proteins (1–391 aa; 5 μg each). *M*r, molecular mass markers. (C) EMSA of the Swi2N derivatives. Linear dsDNA (400 bp) was mixed with increasing concentrations of Swi2N, and the resultant DNA-protein complexes were analyzed by 6% native PAGE. The dsDNA sequences used were *SRE3* (top), *SRE2* (middle), and the region next to *H1* of *mat1* (bottom). (D) EMSA of Swi2N with ssDNA derived from the *H1* homology box (59 nucleotides).

Swi2 localizes to the AT-rich *SRE3* element independently of Swi6 (Jia *et al*., 2004); therefore, we speculated that the AT-hook motifs of Swi2 might have a high affinity for the *SRE3* sequence. To investigate this, we purified Swi2N (1–391 aa) and its AT-hook mutant forms, all of which lacked the C-terminal Sfr1 homology region predicted to possess DNA-binding activity (Figure 3B). In the AT1-A and AT2-A mutants, all residues in the first and second GRP tripeptides, respectively, were mutated to alanine. In the AT12-AA mutant, all residues in the GRP tripeptides in both AT-hooks were mutated to alanine (Figures 3A and 3B).

We examined binding of the AT-hooks to a linear 400 bp dsDNA encompassing the *SRE3* element using EMSA (Figure 3C upper panel). The Swi2N-wild-type (WT) protein efficiently shifted *SRE3* DNA in a protein concentration-dependent manner. Although Swi2N AT1-A and Swi2N AT2-A exhibited only slightly reduced mobility-shifts, Swi2N AT12-AA exhibited drastically reduced band-shifts and could not form stable protein-DNA complexes even at higher protein concentrations. On the other hand, Swi2N-WT did not exhibit a detectable band-shift with ssDNA (Figure 3D). These results clearly indicate that both AT-hooks of Swi2 are involved in dsDNA binding.

Next, we tested the specificity of DNA binding mediated by the AT-hook motifs. An EMSA with Swi2N-WT and *SRE2* or a *mat1* sequence as the target DNA yielded very similar results to those of an EMSA with Swi2N-WT and *SRE3* (Figure 3C middle and lower panels). A similar trend was also observed in an EMSA using the three AT-hook mutant proteins. These results suggest that the AT-hook motifs of Swi2 do not have sequence specificity for *SRE3*.

The results described above were confirmed by performing another EMSA using a non-cognate DNA sequence of bacteriophage ΦX174 (Supplementary Figure S2). When the replicative DNA of ΦX174 (dsDNA) was used both Swi2N-AT1 and Swi2N-AT2 exhibited a similar or slightly reduced band-shift compared with Swi2N-WT, and Swi2N-AT12-AA exhibited a reduced band-shift. When viral DNA (ssDNA) was used, Swi2N-WT and the three AT-hook mutants showed similar weak DNA binding. These results indicate that the AT-hooks of Swi2 are dsDNA-binding motifs that lack specificity for *SRE3*. The finding that Swi2N proteins exhibited a slight shift with viral DNA at higher concentrations may be due to the formation of a secondary structure in viral DNA.

### The C-terminal region of Swi2 homologous to Sfr1 also has non-specific DNA-binding activity

The C-terminal half of Swi2 shares sequence homology with full-length Sfr1 (Akamatsu *et al*., 2003). Sfr1 protein can be roughly divided into two domains: the N- and C-terminal half domains. The N-terminal half is the DNA- and Rad51-binding domain, and the C-terminal half is the Swi5-binding domain (Akamatsu *et al*., 2003; Argunhan *et al*, 2020; Haruta *et al*., 2006; Kuwabara *et al*, 2010; Kuwabara *et al*, 2012). The C-terminal domain cannot be recovered as a soluble protein from *E. coli* cells unless it is co-expressed with Swi5(Haruta *et al*., 2006; Kuwabara *et al*., 2010; Kuwabara *et al*., 2012).

To analyze the biochemical activities of the C-terminal domain of Swi2, we expressed Swi2C (392–722 aa) and Swi2CS (607–722 aa) together with Swi5 in *E. coli* and purified them. For this, we needed to co-express Swi5 as is the case of Sfr1. Purified protein samples are shown in Figure 4A. An EMSA showed modest binding of Swi2C-Swi5 to DNAs cognate to the mating-type region, but without any preference (Figure 4B).

**Figure 4.**
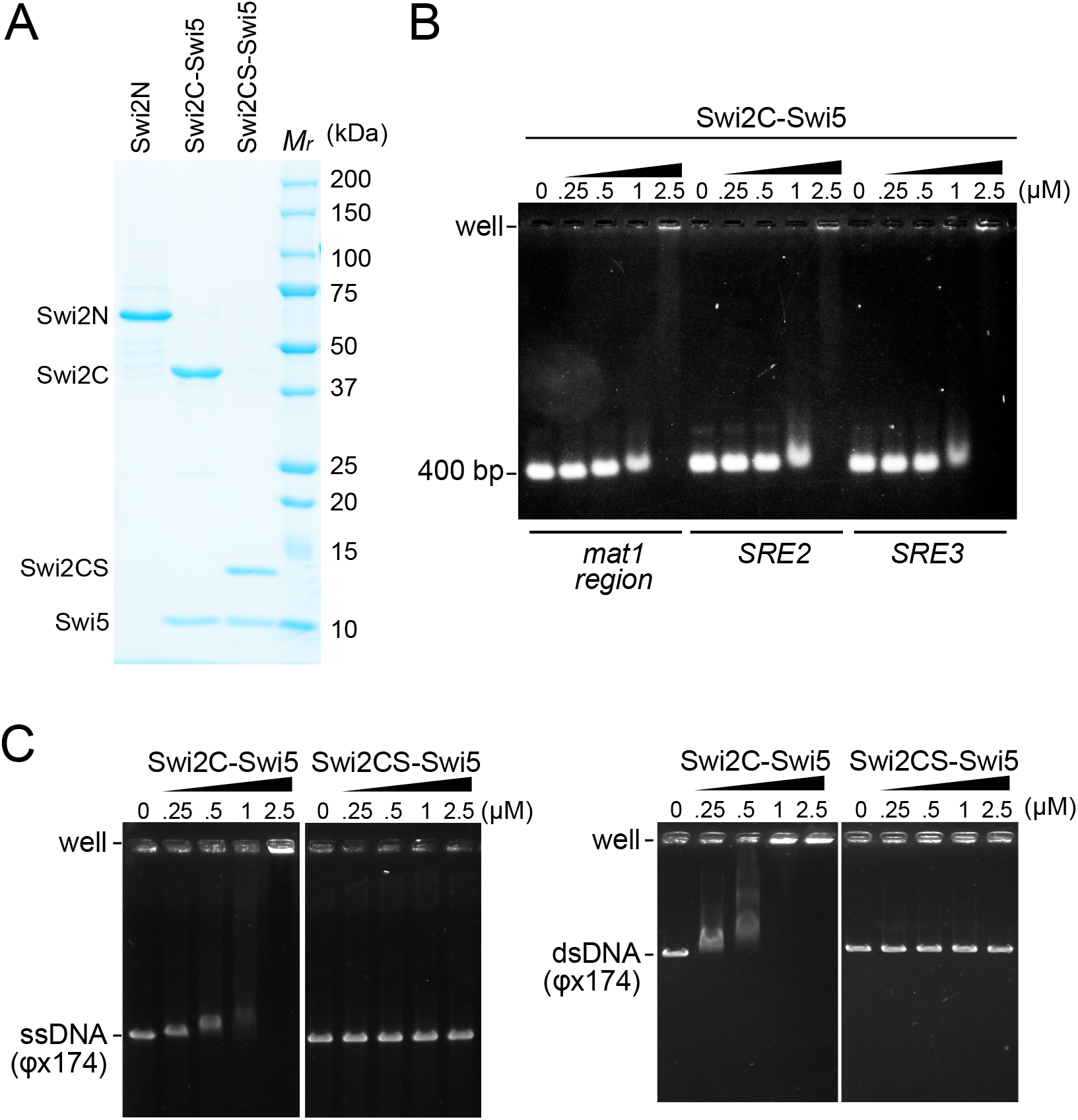
EMSA of the Swi2C-Swi5 and Swi2CS-Swi5 complexes. (A) An SDS-PAGE gel of Swi2N (1–391 aa: 5 μg), the Swi2C (392–722 aa)-Swi5 complex (5 μg), and the Swi2CS (607–722 aa)-Swi5 complex (3 μg). *M*r, molecular mass markers. (B) EMSA of the Swi2C-Swi5 complex. Linear dsDNAs (*mat1, SRE2*, and *SRE3;* 400 bp each) were mixed with increasing concentrations of the Swi2C-Swi5 complex. (C) EMSA of the Swi2C-Swi5 and Swi2CS-Swi5 complexes with a cssDNA (ΦX174 viral DNA) and a circular covalently closed DNA (ΦX174 replicative form DNA).

In contrast to Swi2N, Swi2C-Swi5 showed notable ssDNA binding in an EMSA. However, Swi2CS-Swi5 did not show any detectable binding to ssDNA or dsDNA (Figure 4C), indicating that the region (392–607 aa) is responsible for DNA binding. This region corresponds to the DNA-binding domain of Sfr1 (Argunhan *et al*., 2020; Kuwabara *et al*., 2010).

### The Swi6-binding motif and AT-hook motifs of Swi2 are important for donor choice in MTS

We next examined whether the conserved Swi6-binding residues and DNA-binding AT-hook motifs of Swi2 are involved in MTS. We first performed a classical iodine staining assay (Figure 5A). Among MTS-proficient homothallic *h^90^* cells grown on a minimal medium plate, P and M cells can conjugate. In the assay, spores generated on the plate are darkly stained by iodine vapor. In sharp contrast, MTS-deficient cells are poorly stained because of inefficient spore formation resulting from reduced conjugation. All five *swi2* mutants (Δ: deletion, 5A: Swi6-binding site mutant, AT1-A: first AT-hook mutant, AT2-A: second AT-hook mutant, and AT12-AA: two AT-hooks mutant) formed poorly stained colonies (Figure 5A).

**Figure 5.**
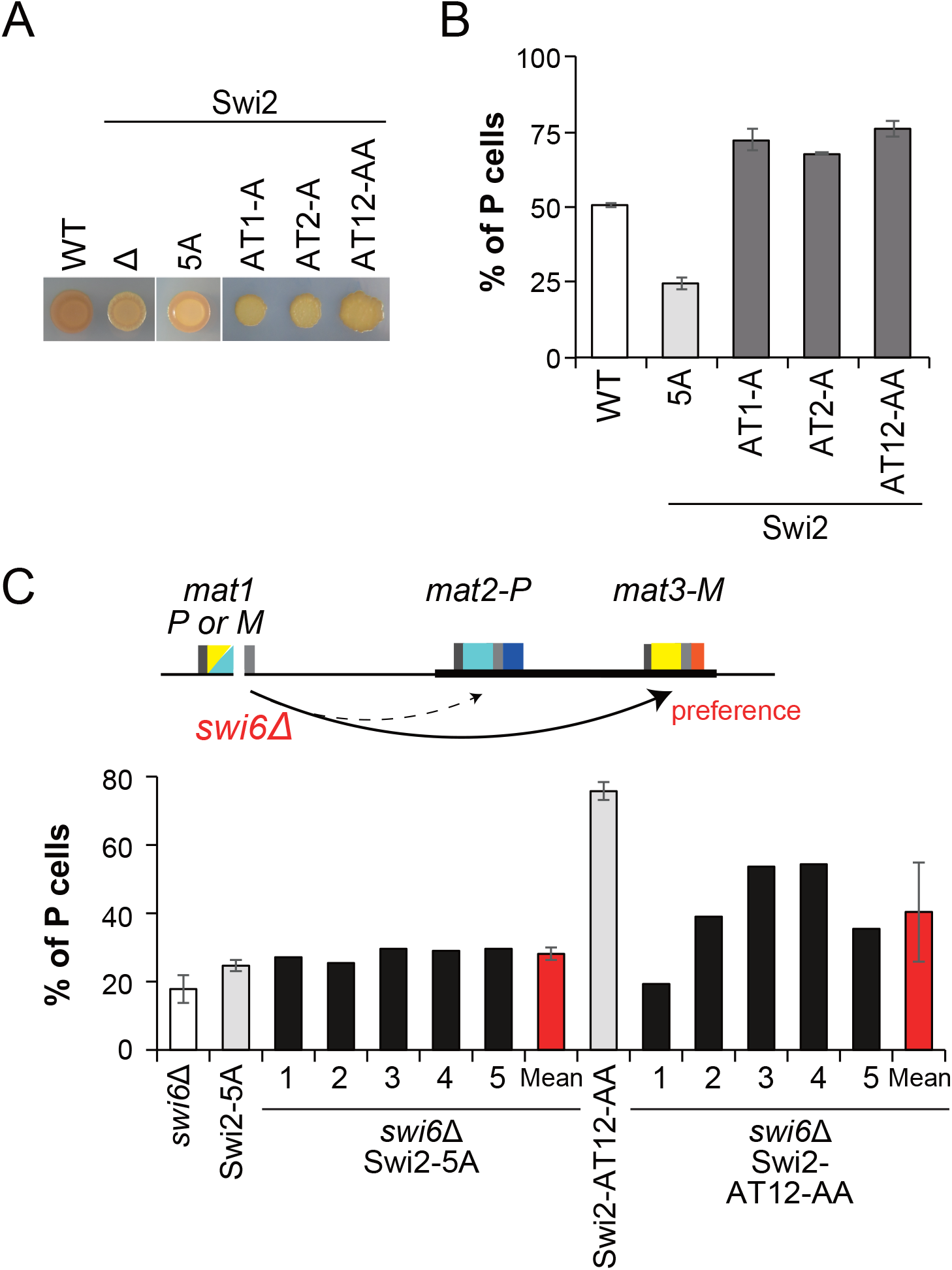
Swi6-binding and AT-hook motifs are required for MTS. (A) Iodine staining of *swi2* mutants. Δ, *swi2* deletion (*swi2Δ);* 5A, Swi6-binding mutant; AT1-A, GRP tripeptide mutant of AT-hook 1; AT2-A, GRP tripeptide mutant of AT-hook 2; and AT12-AA, GRP tripeptide mutant of both AT-hook motifs. (B) The proportion of P cells (%) in the *swi2* mutants was analyzed by multiplex PCR. The graph shows the mean value ± SD for cultures of the WT strain (four cultures of the same clone) or mutants of Swi2-5A (six cultures of independent clones) and AT-hook motifs (three cultures of independent clones). (C) Proportion of P cells (%) for *swi2* mutants in the *swi6*Δ background. Mean values ± SD of the percentage of the P cells of five independent clones of Swi2-5A *swi6*Δ and Swi2-AT12-AA *swi6*Δ.

We next analyzed the P cell population in saturated liquid cultures by multiplex PCR (Figure 5B and Supplementary Figure S3A). WT *h*^90^ cells have a balanced P/M cell ratio. On the other hand, *swi2*Δ cells exhibit clonal differences that cause a bias toward P or M cells (Jakociunas *et al*., 2013). Thus, we analyzed six independent clones of the *swi2-5A* strain and three independent clones of each Swi2 AT-hook mutant strain, *AT1-A, AT2-A*, and *AT12-AA* (Figure 5B and Supplementary Figure S3A). None of these strains exhibited clonal differences, but the Swi6-binding site mutant Swi2-5A and the three AT-hook mutant strains showed an opposite bias in their P/M cell population. The Swi2-5A mutant exhibited a bias toward M cells (25 ± 2% P cells), while each of the AT-hook mutants, *AT1-A, AT2-A*, and *AT12-AA*, exhibited a bias toward P cells (72 ± 3%, 68 ± 0.3%, and 76 ± 3% P cells, respectively) (Figure 5B).

To perform epistasis analysis, we combined the mutations in *swi2* with a deletion of *swi6* (Figure 5C and Supplementary Figure S3B). The proportion of P cells was 18 ± 4% in the single *swi6*Δ mutant. The five independent clones of the *swi2-5A swi6*Δ double mutant strain showed a similar or slightly higher P cell proportion than the *swi2-5A* strain (28 ± 2% P cells). On the other hand, the five independent clones of *swi2-AT12-AA swi6*Δ showed various P cell proportions (31–60% P cells; mean ± standard deviation [SD] of 40 ± 15%).

These results suggest that the MTS defect of the Swi6-binding site mutant of Swi2 (Swi2-5A) is mainly due to a lack of functional interactions with Swi6, leading to a defect in *mat2-P* utilization. Moreover, they suggest that the AT-hook motifs are involved in *mat3-M* utilization and/or inhibition of *mat2-P* utilization. The finding that *swi6*Δ did not completely suppress the bias toward P cells in the *swi2-AT12-AA* mutant also suggests that the AT-hook mutant protein functions, at least partly, independently of Swi6.

### The Swi6-binding and AT-hook motifs are involved in the differential localizations of Swi2 at the two *SRE* elements

The donor choice mechanism is regulated by the cell-type specific *SRE* utilization of Swi2 (Aguilar-Arnal *et al*, 2008; Jakociunas *et al*., 2013; Jia *et al*., 2004; Yu *et al*., 2012). Therefore, we hypothesized that the two interactions of Swi2 with *SRE2* and/or *SRE3* through the Swi6 binding and AT-hook motifs found in this study have a distinct impact on its localization at the *mat* locus. To examine the localization of wild-type and mutant Swi2, we performed a ChIP-qPCR assay using an anti-V5 antibody and Swi2 proteins tagged with 9×V5, in heterothallic strains with a fixed mating-type (Figure 6A). As reported previously (Jia *et al*., 2004), wild-type Swi2 protein abundantly accumulated at *SRE3*, but not at *SRE2*, in P cells, but accumulated at both *SRE2* and *SRE3* in M cells.

**Figure 6.**
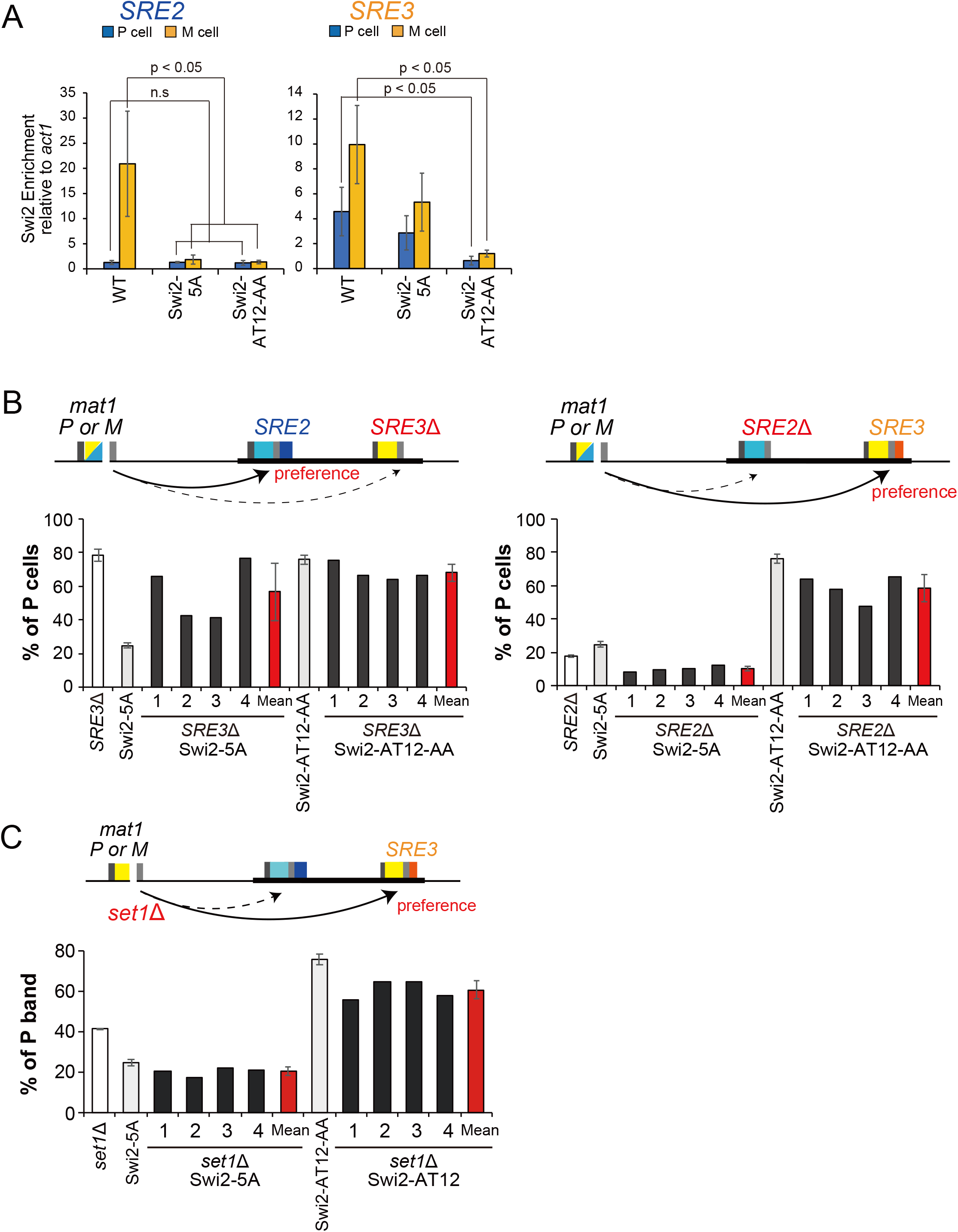
Swi6-binding sites and AT-hook motifs play a crucial role in localization of Swi2 to *SRE* elements. (A) ChIP-qPCR analysis of the Swi2 protein levels relative to that of *act1* in P and M cells. Error bars indicate SD (n=3–5). The two-tailed paired Student’s *t*-test was used to compare the mean values obtained for each mutant with the WT of the same mating type. n.s, not significant. (B) Quantification of the *mat1* contents of each Swi2 mutant strain combined with *SRE3*Δ or *SRE2*Δ by multiplex PCR. Mean values ± SD of the percentage of P cells in four independent clones of strains with mutated *SRE* element (*SRE3*Δ and *SRE2*Δ). (C) Quantification of the *mat1* contents of each Swi2 mutant strain combined with *set1*Δ by multiplex PCR. Mean values ± SD of the percentage of the P cells in four independent clones with *set1*Δ.

In sharp contrast, the Swi2-5A mutant defective in Swi6 binding exhibited deficient localization to *SRE2* in M cells although a low level of association remained. Localization to *SRE2* remained low in P cells, and localization to *SRE3* remained high in both P and M cells. These results are consistent with the previous finding that Swi6 is required for the detection of Swi2 at *SRE2* (Jia *et al*., 2004).

In the case of the AT-hook mutant (*swi2-AT12-AA*), the localization of the mutant Swi2 protein to *SRE2* and *SRE3* was significantly reduced to nearly the basal level in both P and M cells, suggesting that the AT-hook motifs are required for the localization of Swi2 not only to *SRE3* but also to *SRE2*.

### The AT-hook and Swi6-binding motifs are involved in the donor choice mechanism

To examine the impact of Swi2 motif mutation on the donor choice, we measured the P/M cell ratio in *SRE* deletion cells by the multiplex PCR assay (Figure 6B and Supplementary Figure S3C).

In the presence of wild-type Swi2, *SRE3*Δ cultures are predominantly and uniformly of the P mating-type (75–82% P cells; mean ± SD of 78 ± 4%). In contrast, four independent clones of the Swi2-5A mutant in the *SRE3*Δ background showed varied cell-type compositions (41–76% P cells; mean ± SD of 57 ± 17%), thus differing both from *swi2^+^ SRE3*Δ cultures, and from *swi2-5A SRE3^+^* cultures whose bias is uniformly towards M (22–27% P cells; mean ± SD of 25 ± 2%). These phenotypes are compatible with functionally distinct defects in the *swi2-5A* and *SRE3*Δ mutant, whose combination would lead to poor MTS and clonal variation. On the other hand, four independent clones of the Swi2-AT12-AA mutant in the *S-E3*Δ background showed uniform, high proportions of P cells. In spite of a tendency for the proportion of P cells to be slightly lower in the *SRE3*Δ (68 ± 5% P cells) than in the *SRE3^+^* background (76 ± 3% P cells), the strong bias towards P cells in both single and double mutants indicates that Swi2-AT12-AA might act at least in part by reducing the interaction of Swi2 with *SRE3*.

In the *SRE2*Δ background, cultures are predominantly and uniformly of the M mating-type when wild-type Swi2 is present (18 ± 1% P cells). Four independent clones of the Swi2-5A mutant strain in the *SRE2*Δ background had an even stronger bias with a lower proportion of P cells (10 ± 2% P cells) than the *swi2^+^ SRE2*Δ (18 ± 1% P cells) and *swi2-5A SRE2^+^* (25 ± 2% P cells) strains. In sharp contrast, four independent clones of the Swi2-AT12-AA mutant in the *SRE2*Δ background showed a strong bias toward P cells with slightly elevated clonal variation (48–65% P cells; mean ± SD of 58 ± 8% P cells), indicating that they preferentially – albeit somewhat inefficiently – utilize *mat2-P* independently of *SRE2*. For comparison, the Swi2-AT12-AA mutant in the *SRE2^+^* background showed a greater bias toward P cells (76 ± 3% P cells), without detectable clonal variation.

Recently, we reported that two euchromatin factors, Set1C and HULC, inhibit the choice of *mat3-M* as a donor in M cells, probably by affecting chromatin structure at *SRE3* (Esquivel-Chavez *et al*., 2022; Maki *et al*., 2018). We analyzed the possible involvement of these euchromatin factors in the function of Swi2 (Figure 6C and Supplementary Figure S3D). Epistatic analysis showed that the *swi2-5A set1*Δ strain was slightly more biased toward M cells (20 ± 2% P cells) than the *swi2-5A set1^+^* strain (25 ± 2% P cells), while the *swi2^+^ set1*Δ strain had 41 ± 0.3% P cells. On the other hand, the *set1*Δ mutation moderately alleviated the bias toward P cells in the Swi2-AT12-AA strain (reduced from 76 ± 3% to 61 ± 5% P cells) compared to *set1^+^* background, although clonal variation was still observed. These results show that Set1C inhibits the *mat3-M* donor choice even in these Swi2 mutants, independently of the functions of the Swi2 binding and AT-hook motifs.

### The Swi2-Swi5 complex promotes the Rad51-driven DNA strand exchange reaction *in vitro*

The C-terminal region of Swi2 is well-conserved in fission yeasts and shares sequence similarity with Sfr1 protein (Figure 7A and Supplementary Figure S1) (Akamatsu *et al*., 2003). Similar to Swi2, Sfr1 forms a tight complex with Swi5. The Swi5-Sfr1 complex is evolutionarily well-conserved in eukaryotes and promotes the Rad51-driven strand exchange reaction *in vitro* (Haruta *et al*., 2006; Tsai *et al*, 2012). Based on the sequence similarity of Sfr1 and Swi2 (Akamatsu *et al*., 2003), we hypothesized that the Swi2-Swi5 complex would also have strand exchange activity. To test this hypothesis *in vitro*, we purified the Swi2-Swi5 complex from *E. coli* producing Swi2 and Swi5 under the control of the T7 expression system (Figure 7B). In M cells, Swi2 is expressed as a short mRNA (Swi2S: 205–722 aa, as a putative protein) and a full-length mRNA (Yu *et al*., 2012); therefore, we also purified Swi2S as a complex with Swi5 (Figure 7A and B). Neither Swi2 nor Swi2S alone, without Swi5, could be expressed in *E. coli*, as reported previously for Sfr1 (Haruta *et al*., 2006).

**Figure 7.**
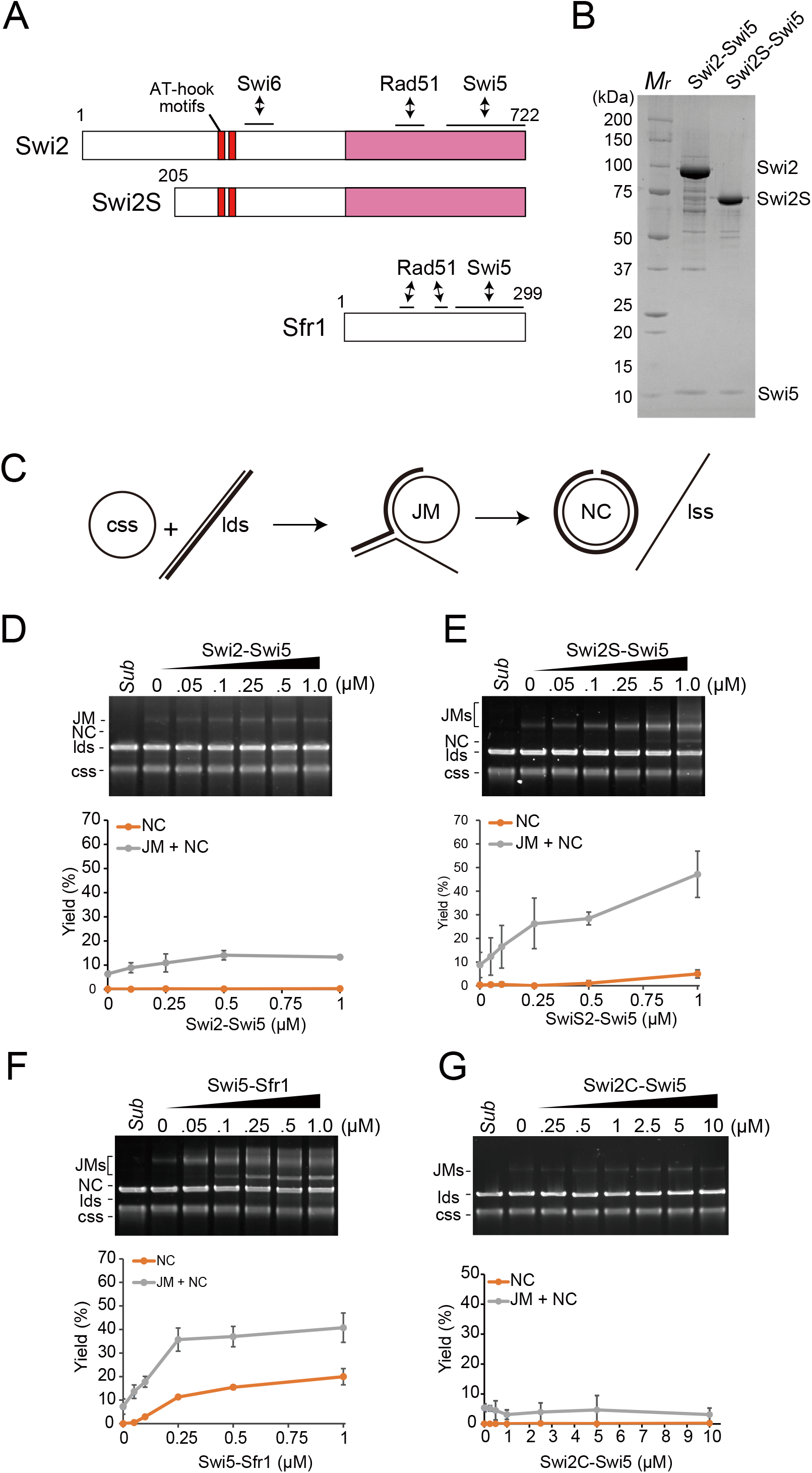
The Swi2-Swi5 complex promotes the Rad51-driven DNA strand exchange reaction. (A) Schematic representation of Swi2 and Swi2S proteins. AT-hook motifs and the interfaces with Swi6 are shown. Swi2S is a putative short form (1–204 aa truncation) of Swi2 expressed in M cells (Yu *et al*., 2012). The C-terminal half region of Swi2 indicated in magenta exhibits sequence similarity with Sfr1 protein. (B) An SDS-PAGE gel of the purified Swi2-Swi5 (8 μg) and Swi2S-Swi5 (5.5 μg) complexes. *M*r, molecular mass markers. (C) Diagram of the three-strand exchange assay. (D–F) Three-strand exchange reactions containing Rad51 (5 μM) and the indicated concentrations of Swi2-Swi5 (D), Swi2S-Swi5 (E), Swi5-Sfr1 (F) or Swi2C-Swi5 (G) were incubated for 2 h at 30°C. DNA was separated by agarose gel electrophoresis to visualize substrates (cssDNA and ldsDNA), intermediates (JM), and products (NC). *Sub*, Substrate. The yields of the NC product (orange) and total yields of the JM intermediate plus NC product (gray) were quantified from the gels. Mean values of three independent experiments ± SD are shown.

We first performed an *in vitro* three-strand DNA exchange assay with φX174 viral cssDNA and linearized double-stranded DNA (ldsDNA) as substrates. The pairing of cssDNA with ldsDNA produces DNA joint molecules (JMs) as intermediates and a nicked circular duplex (NC) as the final product of strand transfer over the length of the substrates (Figure 7C). In Rad51-only reactions, JMs formed (less than 10% yield of the input) and NC was not detected. The addition of the Swi2-Swi5 complex increased JM formation (~14% of the input) (Figure 7D). On the other hand, the addition of the Swi2S-Swi5 complex increased JM and NC formation (~42% and 6% of the input, respectively) (Figure 7E). The addition of the Swi5-Sfr1 complex increased not only JM (~40%) but also NC (~20%) formation (Figure 7F). Importantly, the addition of Swi2C-Swi5, in which Swi2C corresponds to the Sfr1 homology structure (Figure 7A Supplementary Figure S1), did not give any detectable impact on the Rad51-driven strand exchange reaction (Figure 7G). These results suggest that Swi2-Swi5 stimulates Rad51-driven strand-exchange reaction and that not only the C-terminal Sfr1 homology region (392-722 aa) but also half of the long N-terminal (205-392aa) of Swi2 is needed for this stimulation. In addition, this stimulation is suggested to be inhibited by the very N-terminal region (1-204 aa) of Swi2.

Next, we examined whether the Swi2-Swi5 complex promotes Rad51-driven strand invasion into the *H1* homology box *in vitro*. To this end, we employed a D-loop assay using FAM-labeled ssDNA containing 59 bases of the *H1* sequence and scDNA in which the *REIII-mat3-M-H1-SRE3* region was inserted (Figure 8A). In Rad51-only reactions, D-loop formation was undetectable (Figure 8B). Addition of Swi2-Swi5 or Swi2S-Swi5 complex did not markedly increase D-loop formation (less than 2% of the input) (Figure 8C). The Swi5-Sfr1 complex did not promote D-loop formation under the experimental condition (Figure 8C). Similar to Rad51, Rad54 is reportedly essential for MTS (Roseaulin *et al*., 2008).

**Figure 8.**
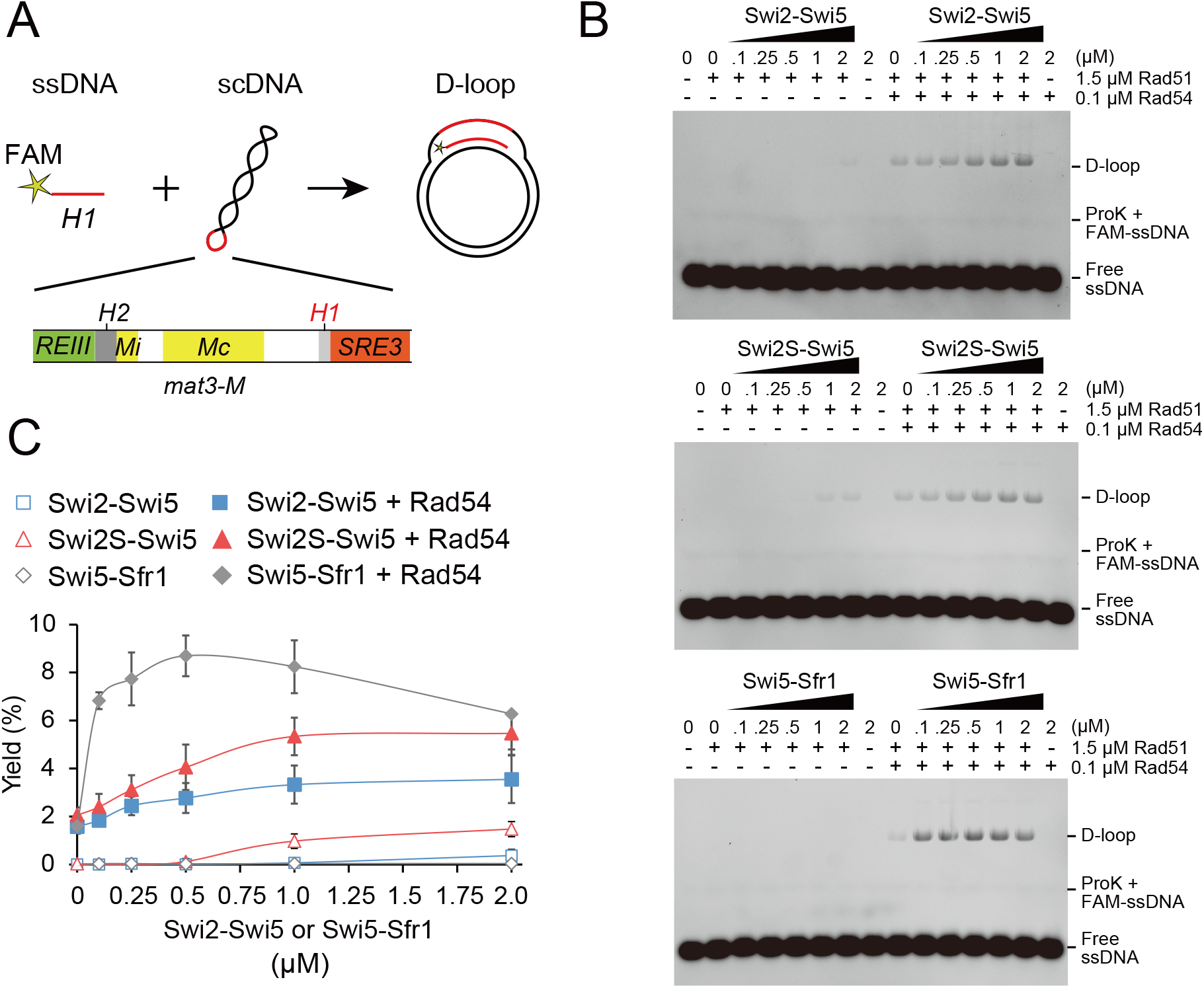
The Swi2-Swi5 complex promotes Rad51-driven D-loop formation. (A) Diagram of the D-loop assay. ssDNA contained the sequence of the *H1* box and was labeled by FAM. scDNA was pBluescript II SK (+) containing ~2300 bp from *REIII* to *SRE3* in the EcoRV site. (B) D-loop reactions containing Rad51 (1.5 μM) and/or Rad54 (0.1 μM) with the indicated concentrations of Swi2-Swi5, Swi2S-Swi5, and Swi5-Sfr1 were incubated for 10 min at 30°C. DNA was separated by agarose gel electrophoresis to visualize substrates. (C) Yields of the D-loop were quantified from the gels (B). The percentage of D-loop signal per total DNA signal was plotted. Mean values of three independent experiments ± SD are shown.

Rad54 proteins from *S. cerevisiae* and human cells promote Rad51-driven D-loop formation *in vitro* (Bugreev *et al*, 2007; Petukhova *et al*, 1998). Therefore, we investigated whether Rad54 stimulates Rad51-driven D-loop formation (Figure 8B). Addition of Rad54 to Rad51-only reactions resulted in very low D-loop formation (~2%) (Figure 8C). D-loop formation was slightly increased by addition of the Swi2-Swi5 complex to the Rad51-Rad54 reaction (~3.5%) and was increased more by addition of the Swi2S-Swi5 complex (~5.5%). D-loop formation greatly increased upon addition of the Swi5-Sfr1 complex (~8.7%). D-loop formation could not be detected when Rad51 was not added to the reactions.

These results indicate that the Swi2-Swi5 complex, independently of Rad54, promotes the Rad51-driven strand exchange reaction *in vitro*. The shorter Swi2S-Swi5 complex has higher activation potential for Rad51-driven strand invasion than the full-length Swi2-Swi5 complex. However, the promotion of Rad51-driven strand exchange by the Swi2S-Swi5 complex is-not as high as that of the Swi5-Sfr1 complex.

### The Swi2-Swi5 complex forms droplets *in vitro*

During the purification steps, we observed that turbidity developed in solutions of Swi2-Swi5 and Swi2S-Swi5 complexes at high protein concentrations, low salt concentrations, and high temperatures. In the presence of 50 and 200 mM NaCl, optical microscopy revealed the presence of clumps in solutions of Swi2-Swi5 complex, and particularly in 50 mM salt solutions containing the Swi2S-Swi5 complex (Figure 9A). By contrast, clumps were not observed in solutions containing the Swi5-Sfr1 complex. We quantified the number of clumps by measuring the turbidity of the solutions at 600 nm (Figure 9B). The turbidity of the solution containing the Swi2-Swi5 complex was higher in the presence of 50 mM NaCl than in the presence of 200 mM NaCl. The salt concentration also influenced the turbidity of solutions containing the Swi2S-Swi5 complex. Consistent with the microscopic observations, the turbidity of the solution containing the Swi2S-Swi5 complex was higher than that of the solution containing the Swi2-Swi5 complex. The turbidity was detected, albeit very low level, for the solution containing Swi2C-Swi5 complex in the presence of 50 mM NaCl. Turbidity was neither detected in solutions containing Swi2N, under either salt conditions nor in solutions containing the Swi5-Sfr1 complex. These results, summarized in Figure 9C, suggest that the molecular characteristics of the Swi2-Swi5 and Swi5-Sfr1 complexes are different. In addition, it is possible that the N-terminus of Swi2 in the Swi2-Swi5 complex mediates self-association, resulting in the formation of droplets.

**Figure 9.**
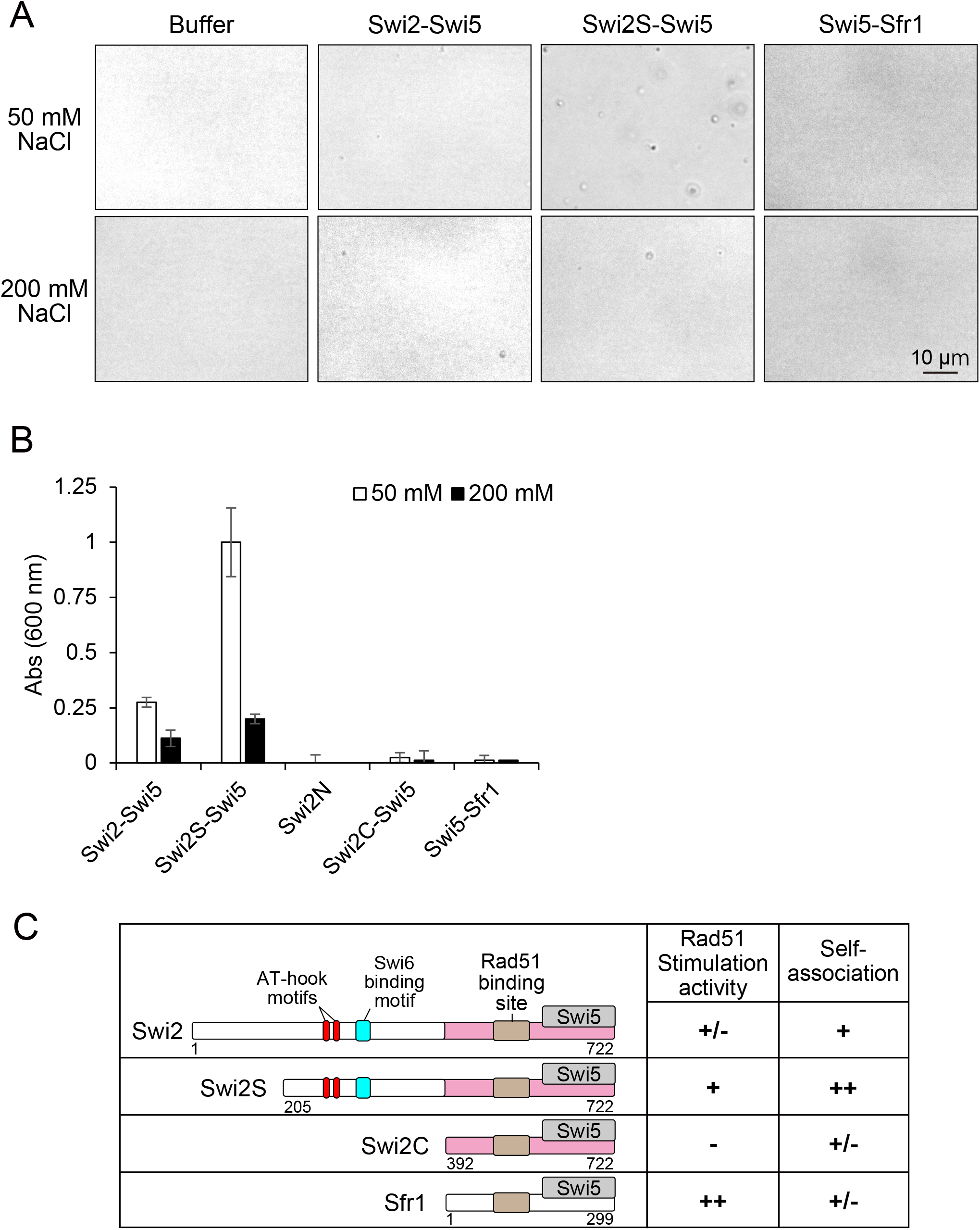
The Swi2-Swi5 and Swi2S-Swi5 complexes form droplets. (A) Optical microscopic images of solutions (10 mM Hepes [pH 7.5] and 0.5 mM DTT) containing 0.5 μM Swi2-Swi5, Swi2S-Swi5, or Swi5-Sfr1 at 25°C and the indicated salt concentrations. (B) The turbidity (*A*600) of solutions containing the indicated proteins and various NaCl concentrations at 25°C was measured using a NanoDrop instrument. The solution was the same as in (A), but each protein concentration was 5 μM. Mean values of three independent experiments ± SD are shown. (C) Summarized properties of purified Swi2-Swi5 and Swi5-Sfr1. Swi2, 1-722 aa; Swi2S, 205-722 aa; Swi2C, 392-722 aa; Sfr1, 1-299 aa. AT-hook motifs are shown in red, the Swi6-binding site in blue, and Rad51-binding site in ochre. The Sfr1 homology region in Swi2 is shown in pink. Stimulation of Rad51-dependent strandexchange activity and self-association are as follows: ++, very strong +, strong; +/-, weak or not detectable; -, not detectable.

## Discussion

The donor choice mechanism in MTS is highly regulated by the MTS factor Swi2-Swi5, chromatin formation, and *cis*-acting *SRE2* and *SRE3* elements (Thon *et al*., 2019). The Swi2-Swi5 complex localizes to *SRE3* independent of Swi6 and to *SRE2* dependent on Swi6, and has been hypothesized to promote the strand exchange reaction for MTS (Jakociunas *et al*., 2013; Jia *et al*., 2004). Here, we identified two functionally important sites in Swi2, the Swi6-binding site, and DNA-binding AT-hook motifs. These two sites were involved in different donor preferences in MTS: the Swi6-binding site was required for the *mat2-P* donor choice, while the AT-hook motifs were required for the *mat3-M* donor choice. In addition, genetic analysis revealed that the binding sites are involved in an *SRE*-mediated function. Moreover, we demonstrated that the Swi2-Swi5 complex promotes Rad51-driven strand invasion *in vitro* and Rad54 further stimulates the activation of strand invasion of Rad51 by the Swi2-Swi5 complex.

### Features of the Swi6-binding site and two AT-hook motifs in Swi2

HP1 was reported to form a platform for various heterochromatin localization proteins that contain a consensus PxVxL sequence as a HP1-binding motif (Smothers & Henikoff, 2000). More recently, the binding consensus was proposed to be a more relaxed Φx(V/P)xΦ sequence (Φ and x indicate a hydrophobic and any amino acid, respectively) (Liu *et al*, 2017). In *S. pombe*, proteins containing Φx(V/P)xΦ, such as Sgo1, Clr3, and Mit1, bind to Swi6 and its paralog Chp2 (Isaac *et al*, 2017; Leopold *et al*, 2019; Yamagishi *et al*, 2008). In Mit1, the xx(I/V)x(I/V) sequence is suggested to be the binding motif (Leopold *et al*., 2019). Swi2 contains two overlapping relaxed consensuses, ^273-^<u>A</u>V<u>V</u>I<u>P</u>^-277^ and ^274-^<u>V</u>VV<u>I</u>P<u>V</u>^-278^ (the underlined amino acids correspond to the consensus sequences), which correspond to the Swi6-binding site identified in this study (Figure 2). Further work is needed to determine which aa exactly are important for Swi6 binding.

We also found two AT-hook motifs near the Swi6-binding motif (Figure 3). The AT-hook motif preferentially binds to the minor groove of A/T-rich DNA (Reeves, 2001). Therefore, because *in vivo* Swi2 constitutively associates with *SRE3*, which is an A/T-rich sequence, we tested whether Swi2 preferentially binds to *SRE3* DNA *in vitro*. However, the results of EMSA demonstrated that the AT-hook motifs bound to the *SRE3* element in basically the same way as they bind to the *SRE2* element. The AT nucleotide contents of the used *mat1* region and *SRE2* and *SRE3* elements were 60%, 75%, and 72%, respectively. The preferential localization of Swi2 to *SRE3 in vivo* may be dependent on other factors such as DNA topology and/or chromatin formation.

Considering that the AT-hook and Swi6-binding motifs are located close to each other in the N-terminal region of Swi2, the configuration of these two motifs may be involved in cooperative and concerted Swi2 bindings to DNA and heterochromatin at *SREs in vivo*, allowing Swi2 to sense the different chromatin contexts in P and M cells.

### Roles of the Swi6-binding and AT-hook motifs of Swi2 in donor designation

The interaction between Swi2 and Swi6 is crucial for *mat2-P* donor choice; therefore, it is well accepted that Swi6-mediated localization of Swi2 to *SRE2* in M cells leads to the selection of *mat2-P* as a donor (Jakociunas *et al*., 2013; Jia *et al*., 2004; Thon *et al*., 2019). In this study, a multiplex PCR assay showed that the Swi6-binding motif mutant *swi2-5A* had a reduced proportion of P cells (25 ± 2% P cells) (Figure 5B), consistent with the proposed model. The ChIP-qPCR assay in M cells showed that accumulation of the Swi2-5A mutant protein was dramatically reduced at *SRE2* but was only slightly decreased at *SRE3*. Moreover, the accumulation of WT and Swi2-5A mutant proteins at *SRE2* and *SRE3* did not significantly differ in P cells (Figure 6A). These results clearly indicate that the interaction with Swi6 is essential for the accumulation of Swi2 at *SRE2* in M cells but is not required for the accumulation of Swi2 at *SRE3* in either cell type.

On the other hand, the AT-hook mutant proteins showed reduced association both with *SRE2* and *SRE3* in M cells and with *SRE3* in P cells (Figure 6A), suggesting that the AT-hook motifs are involved in Swi2 localization to *SRE2* in M cells and to *SRE3* in P cells. Accordingly, the proportion of P cells was higher in the Swi6-binding mutant Swi2-5A (25 ± 2%) and *SRE2*Δ strain (18 ± 1%) than in the double mutant (10 ± 2%; Figure 6B). This result suggests that the *mat2-P* donor choice is governed by partially independent pathways, one involving the Swi2-Swi6 interaction and the other involving *SRE2*. The Swi2-Swi6-interaction-independent and *SRE2*-dependent pathway may involve the AT-hook motifs of Swi2.

We previously showed that Set1C and HULC participate in the *mat2-P* donor choice by acting on *SRE3* (Esquivel-Chavez *et al*., 2022; Maki *et al*., 2018). The current study showed that the double mutant with Swi6-binding and Set1 mutations had a slightly more severe phenotype, while the Set1 mutation slightly suppressed the P cell-bias phenotype of the AT-hook mutant (Figure 6C). This suggests that Set1C and the AT-hook motifs of Swi2 function in different pathways for donor choice during MTS.

Based on the results obtained in the present and previous studies, we propose a model for the donor choice mechanism (Figure 10). In the model, Swi2 in P cells designates the *mat3-M* donor by localizing to *SRE3* via the AT-hook motifs. On the other hand, the *mat2-P* donor choice in M cells involves a complex mechanism. Swi2 localization to *SRE2* via the Swi6 interaction plays a crucial role. In addition, the AT-hook motifs might both foster interaction of Swi2 directly with *SRE2*, and act indirectly through *SRE3* to increase Swi2 occupancy in the whole region. In M cells, we suggest that Set1C also works at *SRE3* to modulate chromatin conformation (Esquivel-Chavez *et al*., 2022; Maki *et al*., 2018). While further work is necessary to elucidate the molecular mechanisms underlying Swi2 localization, our *in vitro* investigations provide new clues for how Swi2 would stimulate recombination when localized at a donor locus and for its cell-type specific properties.

**Figure 10.**
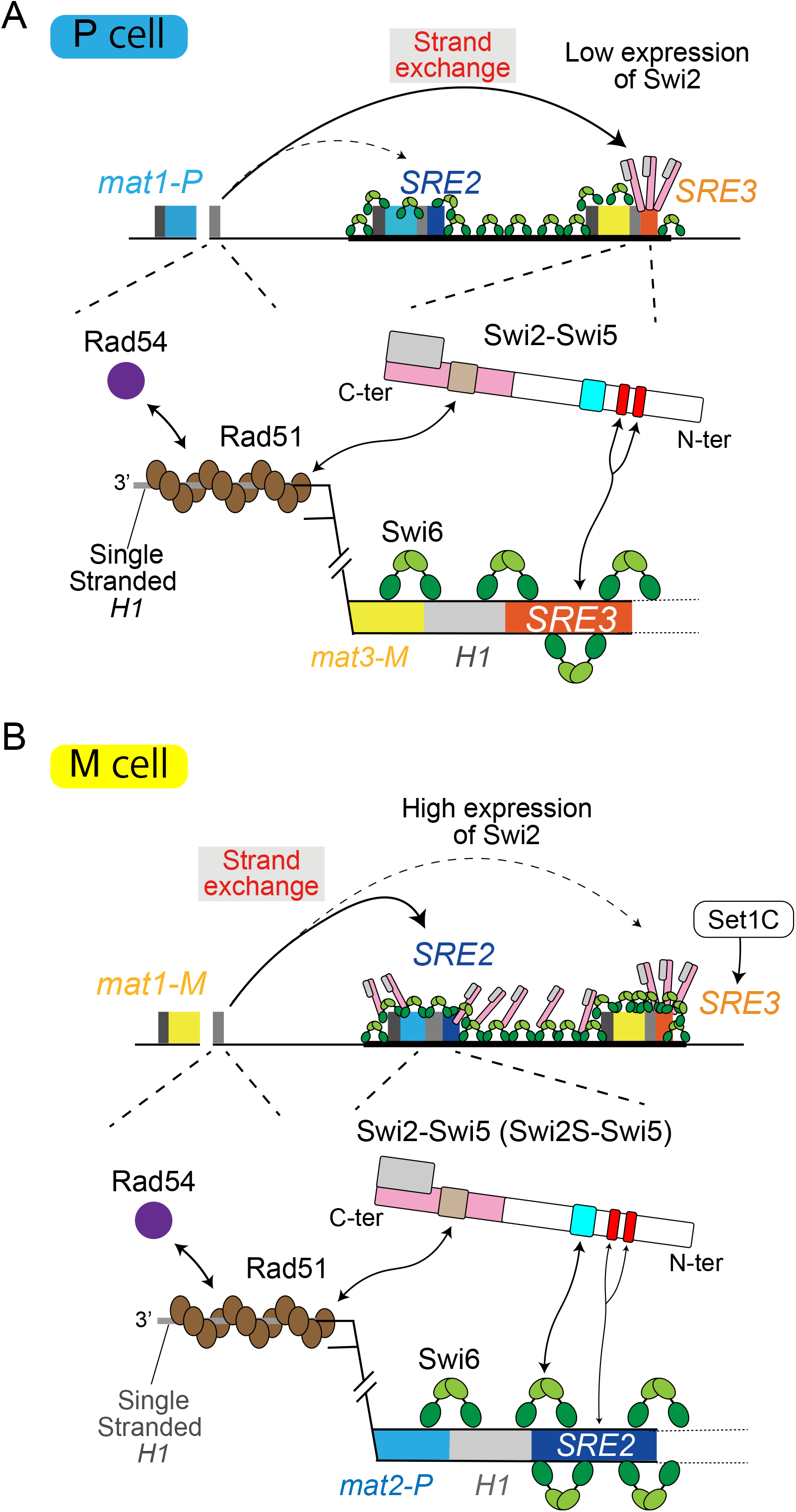
Model for donor choice in MTS. (A) Overall and enlarged views of the *mat* locus in P cells. The Swi2/Swi5 complex localizes to the *SRE3* element via the AT-hook motifs of Swi2 (shown in red). The complex binds to the Rad51 filament formed at *mat1* following resection of *H1* (shown in brown) and stimulates Rad51 strand invasion activity at the *H1* box of the *mat3-M* donor, helped by the mediator of recombination Rad54 (shown in purple). Swi6 is shown as a dimer in green (chromo domain in green and chromo shadow domain in light green). The Swi6-binding motif of Swi2 is in blue, Rad51-binding site in ochre, the Sfr1 homology region in pink and Swi5 is shown in gray. (B) Overall and enlarged views of the *mat* locus in M cells. Swi2, which is in greater abundance in M cells than in P cells (and the shorter Swi2S form of the protein present in M cells) localizes to the *SRE2* element via the Swi6-binding motif, stimulated by the AT-hooks. The Swi2-Swi5 complex on the *SRE2* element binds to Rad51 filament and stimulates strand invasion activity to choose the *mat2-P* donor. The Swi2-Swi5 complex also localizes to the *SRE3* element via the AT-hook motifs; however, Set1C inhibits *mat3-M* donor choice at the *SRE3* element.

### Effect of the Swi2-Swi5 complex on strand exchange driven by Rad51

We demonstrated that the Swi2-Swi5 complex promotes Rad51-driven strand exchange *in vitro* (Figure 7), suggesting that it can directly promote Rad51-driven strand invasion and subsequent strand exchange at the *mat* locus. Addition of Rad54 protein further stimulated the strand invasion activity of Rad51 (Figure 8). Rad54 physically interacts with Rad51 and is involved in DSB repair at the *mat* locus (Afshar *et al*, 2021; Roseaulin *et al*., 2008). We propose a model in which Rad51 and Rad54 bind to ssDNA generated from a one-ended DSB next to the *H1* box of the *mat1* cassette. This occurs in cells of both mating types (Figure 10). The Swi2-Swi5 complex localizes to the *mat* locus without a DSB and is therefore assumed to localize to this locus before a DSB forms and then to recruit the Rad51 filament (Thon *et al*., 2019).

The N-terminal half of Swi2 (1-391aa) in addition to the Sfr1 homology region (392-722aa) plays also an important role in stimulation of Rad51-driven strand-exchange activity (Figure 7). Interestingly, the N-terminal truncated Swi2S-Swi5 complex (lacking 1–204 aa of Swi2) had higher stimulation activity than the full-length Swi2-Swi5 complex (Figures 7 and 8) and the Swi2S-Swi5 complex formed more clumps in solution than the full-length Swi2-Swi5 complex (Figure 9). This suggests that a portion of Swi2 (205-391aa) within the N-terminal half self-associates to facilitate clump formation by the complex and that the very N-terminus (1–204 aa) inhibits both stimulations of Rad51 activity and self-association. Although in this study we could not elicit a rigid conclusion that the observed clump formation was related to liquidliquid phase separation, the self-association of Swi2-Swi5 and Rad51 stimulation by the complex were closely related to each other. Swi2S is assumed to be translated in M cells (Yu *et al*., 2012). The different characteristics of the Swi2-Swi5 and Swi2S-Swi5 complexes may affect *mat2-P* donor choice in M cells (Figure 10).

In this study, we found that the functions of the Swi2-Swi5 complex in donor directionality can be separated between the Swi6-binding motif and the DNA-binding AT-hook motifs of Swi2. Moreover, we provide the first biochemical evidence that the Swi2-Swi5 complex stimulates Rad51-driven strand exchange activity *in vitro*. The uncovered properties of Swi2 - to recognize the chromatin state, bind DNA and stimulate DNA recombination - are central to accurately executing MTS through the spatiotemporal regulation of donor choice. These properties provide a rare glimpse into events occurring at donor sites rather than the more studied events occurring at sites of DNA damage.

## Materials and Methods

### Yeast strains and strain constructions and manipulations

The *S. pombe* strains used in this study are described in Supplementary Table S1. Standard techniques were used to cultivate, sporulate, cross, and genetically manipulate *S. pombe* (Ekwall & Thon, 2017). Strains were generated by transformations or genetic crosses. The *swi2* mutant strains (TM366, TM511, TM519, and TM527) were generated from EM20 by the CRISPR/Cas9 method (Esquivel-Chavez *et al*., 2022). EM20 was transformed with a gRNA expression plasmid (pEM59 derivative), a Cas9 expression plasmid, and donor template DNA for HR. The target sequences were 5’-TCGATGCTGTAGTCATTC-3’ for Swi2-5A, 5’-GAGTGAAGCATAGGGCAGGA-3’ for Swi2-AT1 and -AT12-AA, and 5’-GAAGGATTGCCAAGAAGAAG-3’ for Swi2-AT2-A and -AT12-AA, each of which was inserted into the BbsI site of pEM59. The HR donor templates for mutagenesis were amplified from the *swi2^+^* plasmid (pGADT7-Swi2-5A and pET-SUMO-Swi2N-AT1-A, -Swi2N-AT2-A, and -Swi2N-AT12-AA) using primer sets (Swi2-5A: 5’-GGGGAAGACCTACAGGCTGG-3’ and 5’-CAGGGAAGTCTTCATCCTCTCTTG-3’; and Swi2-AT mutants: 5’-GGTTGGAACCAGCTGTGTG-3’ and 5’-TGAGGTAACGAGGGCTCTG-3’).

### Iodine staining assay

The efficiency of MTS was estimated by iodine staining as described previously (Thon & Klar, 1993). Cells were streaked on MSA (minimal sporulating agar) plates (Petersen & Russell, 2016)and grown at 26°C for 3–4 days. Plates on which colonies were grown were exposed to iodine vapor for ~10 min at room temperature.

### Multiplex PCR assay

Sample preparation and multiplex PCR were performed as previously described (Maki *et al*., 2018). Isolated colonies of the strains were cultured in 2 mL YE5S liquid at 30°C until saturation. Each culture was harvested and genomic DNA was extracted using a Dr. GenTLE (from Yeast) High Recovery Kit (9082, Takara Bio). The genomic DNA concentration was quantified using the Quantifluor® ONE dsDNA Dye System (E4971, Promega). Genomic DNA (5–20 ng) was added to the PCR reaction mixture (total 20 μL). The common forward primer FAM-MT1 (5’-AAATAGTGGGTTAGCCGTGAAAGG-3’), the *mat1-P-*specific reverse primer MP1 (5’-ATCTATCAGGAGATTGGGCAGGTG-3’), and the *mat1-M-*specific reverse primer MM1 (5’-GGGAACCCGCTGATAATTCTTGG-3’) were used at concentrations of 400, 200, and 200 nM, respectively. The 5’ end of FAM-MT1 was labeled with 6-carboxyfluorescein (FAM). To reduce non-specific PCR products, 400 nM heat-stable RecA protein from *Thermus thermophiles* and 400 μM ATP were included in the PCR reaction buffer (10 mM Tris-HCl [pH 8.3], 50 mM KCl, and 2.5 mM MgCl_2_) (Shigemori *et al*, 2005). The amplification program was 2 min at 94°C, followed by 27 cycles of 30 sec at 94°C, 30 sec at 55°C, and 1 min at 72°C, and a final extension of 5 min at 72°C. The PCR fragments corresponding to the *mat1-P* and *mat1-M* alleles were resolved on 5% polyacrylamide gels. Fluorescence was detected and quantified using Typhoon FLA9500 (Cytiva) and ImageQuant (Cytiva).

### Yeast two-hybrid assay

The yeast two-hybrid assay was performed as described previously (Akamatsu *et al*., 2003). pGADT7 plasmids were generated using pGADT7-Swi2 full-length from our laboratory stock by inverse PCR or the quick-change method using the primer sets listed in Supplementary Table S2.

### Protein purification

The *swi2N* (1–391 aa), *swi2^+^* (1–722 aa), *swi2S* (205–722 aa), *swi2C* (392–722 aa), and *swi2CS* (607–722 aa) genes were cloned into a His-SUMO fusion protein expression vector (Kim *et al*, 2018) using an In-Fusion Cloning Kit (639648, Takara Bio). The glycine-arginine-proline (GRP) tripeptides in the AT-hooks of Swi2N were mutated by the quick-change method. The DNA fragments for cloning were amplified using PrimeStar Max DNA Polymerase (R045A, Takara Bio) and the primer sets listed in Supplementary Table S2. Each His-SUMO-Swi2N protein was expressed in *E. coli* BL21-CodonPlus (DE3) and purified as follows. Cells carrying the Swi2N expression plasmid were grown at 37°C until OD600 reached 0.5 in LB medium containing ampicillin. IPTG was added to a final concentration of 0.2 mM, and the culture was further incubated for ~16 h at 18°C. The cells were collected by centrifugation and the pellet was suspended in R buffer (20 mM Tris-HCl [pH 8.0], 1 mM EDTA, 1 mM DTT, and 5% glycerol) containing 200 mM NaCl. The cells were disrupted by sonication and centrifuged at 40,000 × g for 30 min. The supernatant was loaded onto a Ni-NTA Superflow Cartridge (30761, Qiagen). The cartridge was washed with 100 mL R buffer containing 200 mM NaCl. Proteins were eluted with R buffer containing 300 mM imidazole and 200 mM NaCl. The sample was cleaved by His-Ulp1 protein overnight at 4°C in dialysis buffer 1 (20 mM Tris-HCl [pH 7.5], 100 mM NaCl, 5% glycerol, and 1 mM DTT). The flow-through sample of Ni--NTA Superflow Cartridge was loaded onto a 5 mL HiTrap Heparin column (17040701, Cytiva). Proteins were eluted with a linear gradient of 0.1–0.6 M NaCl. Peak fractions of Swi2N protein appeared at 250 mM NaCl. The fractions were diluted three times with R buffer and loaded onto a Resource Q column (17117701, Cytiva). Swi2N eluted at ~250 mM NaCl. The eluted Swi2N was dialyzed against R buffer containing 200 mM NaCl, and concentrated using an Amicon Ultra-4 (30 k) centrifugal filter (UFC803008, Merck Millipore).

Swi2, Swi2S, Swi2C, and Swi2CS proteins were purified in complex with Swi5 (Swi2-Swi5, Swi2S-Swi5, Swi2C-Swi5, and Swi2CS-Swi5). BL21-CodonPlus (DE3) carrying the His-SUMO-Swi2, -Swi2S, -Swi2C, or -Swi2CS expression plasmid and the Swi5 expression plasmid derivative of pET28a was grown at 37°C until OD600 reached 0.5 in LB medium containing ampicillin and kanamycin. IPTG was added to a final concentration of 0.2 mM, and the culture was further incubated for ~16 h at 18°C. The cells were collected by centrifugation and the pellet was suspended in R buffer containing 200 mM NaCl. The cells were disrupted by sonication and centrifuged at 40,000 × g for 30 min. The supernatant was loaded onto a Ni-NTA Superflow Cartridge (30761, Qiagen). The cartridge was washed with 100 mL R buffer containing 200 mM NaCl. Proteins were eluted with R buffer containing 300 mM imidazole and 200 mM NaCl. The eluted proteins were cleaved by His-Ulp1 protein at 4°C for 4 h to overnight in dialysis buffer2 (20 mM Tris-HCl [pH 7.5], 150 mM NaCl, 5% glycerol, and 1 mM DTT). After cleavage of the SUMO tag, each protein complex underwent a different purification procedure. The Swi2-Swi5 sample was applied to a 5 mL HiTrap Q column (17115401, Cytiva). Proteins were eluted with a linear gradient (100 mL of 0.2–0.6 M NaCl). Peak fractions were diluted with two volumes of R buffer and loaded onto a 5 mL HiTrap SP column (17115201, Cytiva). The Swi2-Swi5 complex was eluted with a linear gradient (100 mL of 0.15–0.55 M NaCl). Peak fractions were diluted with an equal volume of R buffer. The sample was loaded onto a 5 mL HiTrap Heparin column (17040701, Cytiva). The Swi2-Swi5 complex was eluted with a linear gradient (30 mL 0.25–0.65 M NaCl). The peak fraction was applied onto a Superdex 200 10/300 GL gel filtration column (17517501, Cytiva) in R buffer containing 200 mM NaCl. Peak fractions containing the Swi2-Swi5 complex were concentrated using an Amicon Ultra-4 (50 k) centrifugal filter (UFC805008, Merck Millipore). After cleavage, the Swi2S-Swi5 complex was applied to a 5 mL HiTrap Q column and washed with R buffer containing 300 mM NaCl. The flow-through of the HiTrap Q column (17115401, Cytiva) was loaded onto a 5 mL HiTrap SP column (17115201, Cytiva). The Swi2S-Swi5 complex was eluted with a linear gradient (100 mL of 0.2–0.6 M NaCl). Peak fractions were diluted with an equal volume of R buffer and loaded onto a 5 mL HiTrap Heparin column (17040701, Cytiva). The Swi2S-Swi5 complex was eluted with a linear gradient (30 mL 0.2–0.8 M NaCl). The peak fraction was developed in a Superdex 200 10/300 GL gel filtration column (17517501, Cytiva) in R buffer containing 200 mM NaCl. The Swi2S-Swi5 complex was concentrated using an Amicon Ultra-4 (50 k) centrifugal filter (UFC805008, Merck Millipore), applied to a 5 mL HiTrap Heparin column, and washed with R buffer containing 150 mM NaCl. The flow-through of the HiTrap Heparin column (17040701, Cytiva) was loaded onto a 5 mL HiTrap SP column (17115201, Cytiva). The Swi2C-Swi5 complex was eluted with a linear gradient (100 mL of 0.1–0.5 M NaCl). Peak fractions were diluted with four volumes of R buffer. The sample was loaded onto a 5 mL HiTrap Q column (17115401, Cytiva). The Swi2C-Swi5 complex was eluted with a linear gradient (50 mL of 0–0.5 M NaCl). The peak fraction was developed in a Superdex 200 10/300 GL gel filtration column (17517501, Cytiva) in R buffer containing 200 mM NaCl. The Swi2C-Swi5 complex was concentrated using an Amicon Ultra-4 (30 k) centrifugal filter (UFC803008, Merck Millipore). After cleavage, the Swi2CS-Swi5 complex was diluted with four volumes of R buffer. The sample was loaded onto a 5 mL HiTrap Q column (17115401, Cytiva). The Swi2CS-Swi5 complex was eluted with a linear gradient (100 mL of 0–0.4 M NaCl). The peak fraction was developed in a Superdex 200 10/300 GL gel filtration column (17517501, Cytiva) in R buffer containing 200 mM NaCl. The Swi2CS-Swi5 complex was concentrated using an Amicon Ultra-4 (10 k) centrifugal filter (UFC801008, Merck Millipore).

*S. pombe* Rad51, Swi5-Sfr1, Swi5-Sfr1C, and RPA proteins were purified as described previously (Haruta *et al*., 2006; Kuwabara *et al*., 2012). *S. pombe* Rad54 protein was purified as described by Afshar *et al*. (Afshar *et al*., 2021) with the following modifications. Rad54 was expressed in a 4 L culture of *E. coli* BL21 (DE3) Star cells carrying the T7-based plasmid pBA110 (pET15b-6xHis-Fh8-PreScission-1xFLAG-*rad54^+^*) as described previously (Afshar *et al*., 2021). The cells were collected by centrifugation and the pellet was suspended in 120 mL R buffer containing 200 mM NaCl. The cells were disrupted by sonication and centrifuged at 40,000 × g for 30 min. The supernatant was loaded onto a 10 mL Ni-NTA Superflow cartridge (connected two 5 mL cartridges) (30761, Qiagen). The cartridge was washed with 100 mL R buffer containing 200 mM NaCl. Proteins were eluted with R buffer containing 300 mM imidazole and 200 mM NaCl. The sample was cleaved by PreScission Protease (27084301, Cytiva) overnight at 4°C in dialysis buffer 1. The protein sample was applied to a 5 mL HiTrap Q column (17115401, Cytiva). The flow-through of the HiTrap Q column was loaded onto a 5 mL HiTrap SP column (17115201, Cytiva). Proteins were eluted with a linear gradient (100 mL of 0.2–0.6 M NaCl). Peak fractions were diluted with two volumes of R buffer. The sample was loaded onto a 5 mL HiTrap Heparin column (17040701, Cytiva). Proteins were eluted with a linear gradient (50 mL of 0.2–0.6 M NaCl). Rad54 was concentrated using an Amicon Ultra-4 (50 k) centrifugal filter (UFC805008, Merck Millipore).

### Electrophoresis mobility shift assay (EMSA)

The target DNA fragments *mat1, SRE2*, and *SRE3* (400 bp each) were amplified by PCR using a laboratory stock plasmid containing the *mat* locus as a template and primer sets (*SRE3:* 5’-TTATCCAAATATGTTTGTTTGGCCGAT-3’ and 5’-GGGTAAGAAGAACTTTTATTTATTTATTTGCC-3’; *SRE2:* 5’-GTTTGTGATTATGCTGTTCAGCATTG-3’ and 5’-GCGAAGCATATTTCTTGCTAATCTTTTG-3’; and *mat1* region: 5’-TTTCCAATTATGCTGTTCGTGTCATTC-3’ and 5’-AGAATGCTCTATGGTTGAGGAAGT-3’). The 5’ end FAM-labeled ssDNA of the *H1* box (59 nucleotides) was purchased from Eurofin Genomics. One of the target DNAs (5 ng/mL final concentration) and Swi2N protein were mixed in buffer (20 mM Tris-HCl [pH7.5], 100 mM NaCl, 5% glycerol, 1 mM DTT) and incubated at room temperature for 10 min. The reactions were analyzed by 1% agarose gel electrophoresis at 50 V for 90 min with TAE buffer. Doublestranded DNA (dsDNA) was stained with GelRed (41002, Biotium) and detected using a Typhoon FLA9500 scanner (Cytiva).

### Chromatin immunoprecipitation (ChIP)-quantitative PCR (qPCR) assay

ChIP was performed essentially as described previously (Esquivel-Chavez *et al*., 2022). EMM2 medium (50 mL) containing 0.1 g/l each of leucine, adenine, histidine, uracil, and arginine was used for cell culture. The cultures were grown to a density of 1.0 × 10^7^ cells/mL at 30°C and then shifted to 18°C for 2 h. A total of 5.0 × 10^8^ cells were cross-linked with 1% formaldehyde for 15 min at 25°C and then incubated in 125 mM glycine for 5 min. Cross-linked cell lysates were solubilized using a multi-beads shocker (Yasui Kikai) and sonicated using a Bioruptor UCD-200 system (Diagenode). The sheared samples were centrifuged at 20,000 g for 10 min at 4°C. The supernatants were incubated with 30 μL Dynabeads Protein A (DB10002, Thermo Fisher) preloaded with 2.5 μL anti-V5 antibody (015-23594, FUJIFILM Wako) for 6 h at 4°C. After washing the beads, materials coprecipitated with the beads were eluted with elution buffer (50 mM Tris-HCl [pH 7.6], 10 mM EDTA, and 1% SDS) for 20 min at 65°C. The eluates were incubated overnight at 65°C to reverse cross-links and then treated with 10 μg/mL RNase A for 1 h at 37°C followed by 20 μg/mL proteinase K for 3 h at 50°C. DNA was purified with a MonoFas DNA Purification Kit I (A01-0004, ANIMOS). qPCR was performed with SYBR Premix Ex Taq II (RR820A, TaKaRa Bio) or TB Green Premix DimerEraser (RR091A, Takara Bio) on a Mx3000P qPCR system (Agilent).

### Three-strand DNA exchange assay

The three-strand assay was performed with reference to a previous study (Ito *et al*, 2020). In total, 5 μM Rad51 and 10 μM (nucleotide) φX174 circular single-stranded DNA (ccsDNA) (N3023L, NEB) were mixed in reaction buffer (25 mM Tris-HCl [pH 7.5], 30 mM KCl, 50 mM NaCl, 2 mM MgCl_2_, 1 mM DTT, 5% glycerol, 2 mM ATP, 8 mM creatine phosphate, and 8 U/mL creatine kinase), and the mixture was incubated at 30°C. After 5 min, the indicated concentration of Swi2-Swi5 or Swi5-Sfr1 was added and incubated for 5 min, followed by addition of 1 μM RPA. After 10 min, 10 μM (nucleotide) linear φX174 dsDNA (N3021L, NEB) was added to the mixture to initiate the three-strand exchange reaction. After 120 min, 200 μg psoralen was added and the reaction mixtures were exposed to ultraviolet (UV) light to crosslink DNA. After DNA cross-linking, 1.8 μl reaction stop solution containing 5.3% SDS and 6.6 mg/mL proteinase K (9034, Takara) was added and the sample was incubated for 60 min at 37°C. Substrates and reaction products were separated by agarose gel electrophoresis and the gel was stained with SYBR-Gold (S11494, Thermo Fisher).

### D-loop assay

A super-coiled plasmid DNA of pBluescript II SK containing *mat3* sequences (*REIII-mat3-H1-SRE3*), which was first prepared by using a Qiagen Plasmid Maxi Kit (12963, Qiagen) and further purified by CsCl equilibrium ultracentrifugation with ethidium bromide (Sambrook *et al*, 1989) was used as a substrate super-coiled DNA (scDNA).

The 5’ end FAM-labeled ssDNA of the *H1* box (59 bases) was purchased from Eurofin Genomics. All components used in the reactions are described in terms of their final concentrations. DNA concentrations are described in terms of total nucleotides. Reactions were performed in 30 mM Tris-HCl [pH 7.5], 40 mM NaCl, 20 mM KCl, 5 mM MgCl_2_, 2% (w/v) glycerol, 2 mM ATP, and 1 mM DTT. Rad51 (1.5 μM) was mixed with FAM-labeled *H1* ssDNA (FAM-H1) (30 nM) and incubated for 5 min at 30°C. A mixture of scDNA (30 nM) and Swi2-Swi5 or Swi5-Sfr1 (concentration described in the figure) was pre-incubated for 15 min at room temperature and then added to the mixture containing Rad51 and ssDNA to initiate D-loop reaction. After incubation for 10 min at 30°C, the reaction was terminated by adding one-fifth of the volume of stop solution (1% (w/v) SDS and 1 mg/mL protease K). The product was separated by 1% agarose gel electrophoresis at 50 V for 90 min with TAE buffer. FAM-H1 was detected and quantified using Typhoon FLA9500 (Cytiva) and ImageQuant (Cytiva).

## Data Availability

The data of this study are available within the article and/or its supplementary data, or available upon reasonable request.

## Acknowledgments

We are grateful to members of the Iwasaki Laboratory, Yumiko Kurokawa, Yasuto Murayama, and James E. Haber for discussions. We thank the National BioResource Project for providing strains. We thank the Biomaterials Analysis Division, Open Facility Center, Tokyo Institute of Technology for sequencing analysis. H. I was supported by Grants-in-Aid for Scientific Research (A) (JP18H03985 and JP22H00404) from the Japan Society for the Promotion of Science (JSPS) and G. T. was supported by grant NNF19OC0058686 from the Novo Nordisk Foundation.

## Author contributions

**Takahisa Maki:** Data curation; formal analysis; validation; investigation; methodology; writing – original draft; writing – review and editing. **Geneviève Thon:** Conceptualization; supervision; funding acquisition; writing – review and editing. **Hiroshi Iwasaki:** Conceptualization; supervision; funding acquisition; investigation; project administration; writing – review and editing.

## Supplementary Figures

**Supplementary Figure S1.**
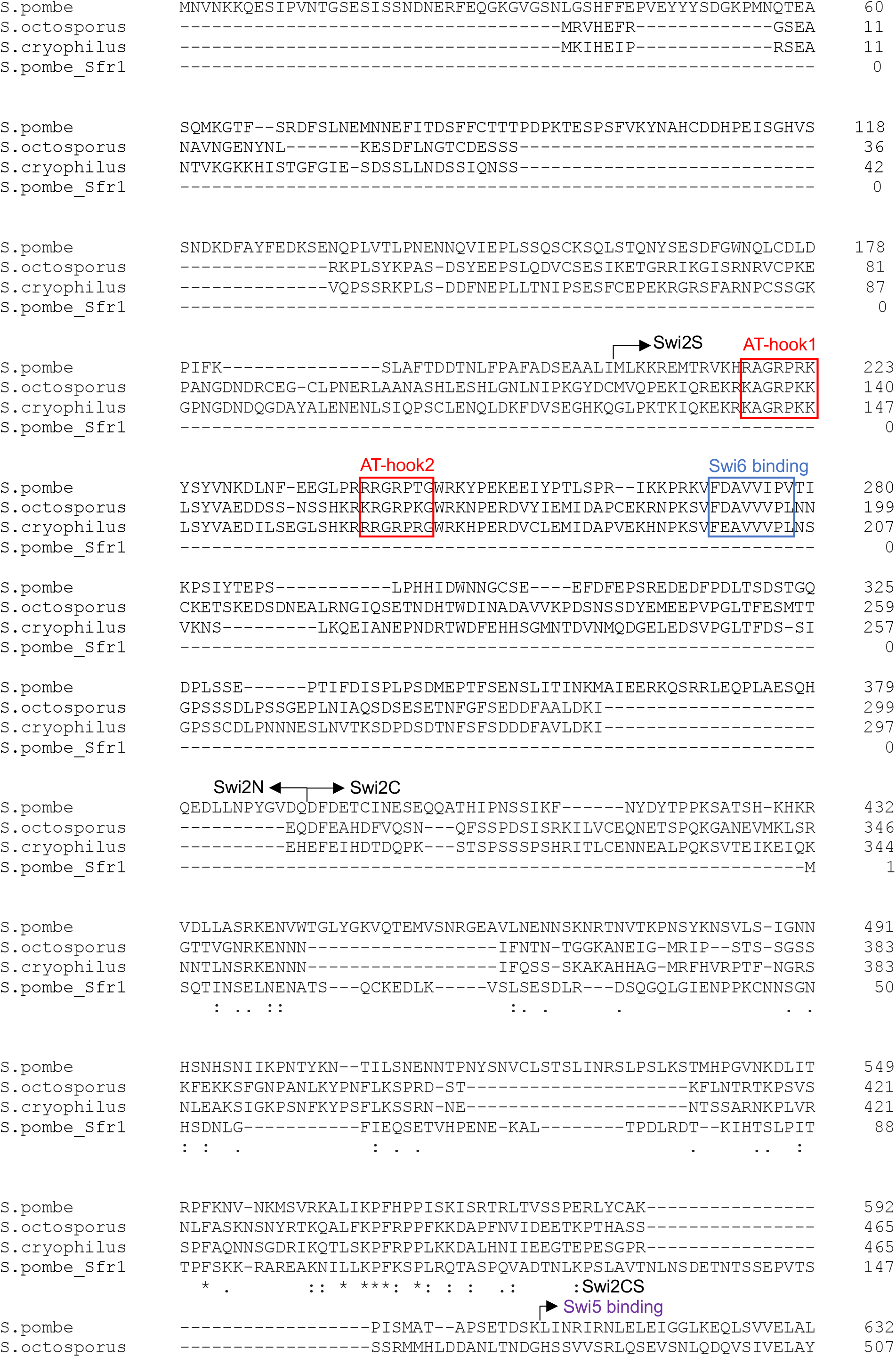
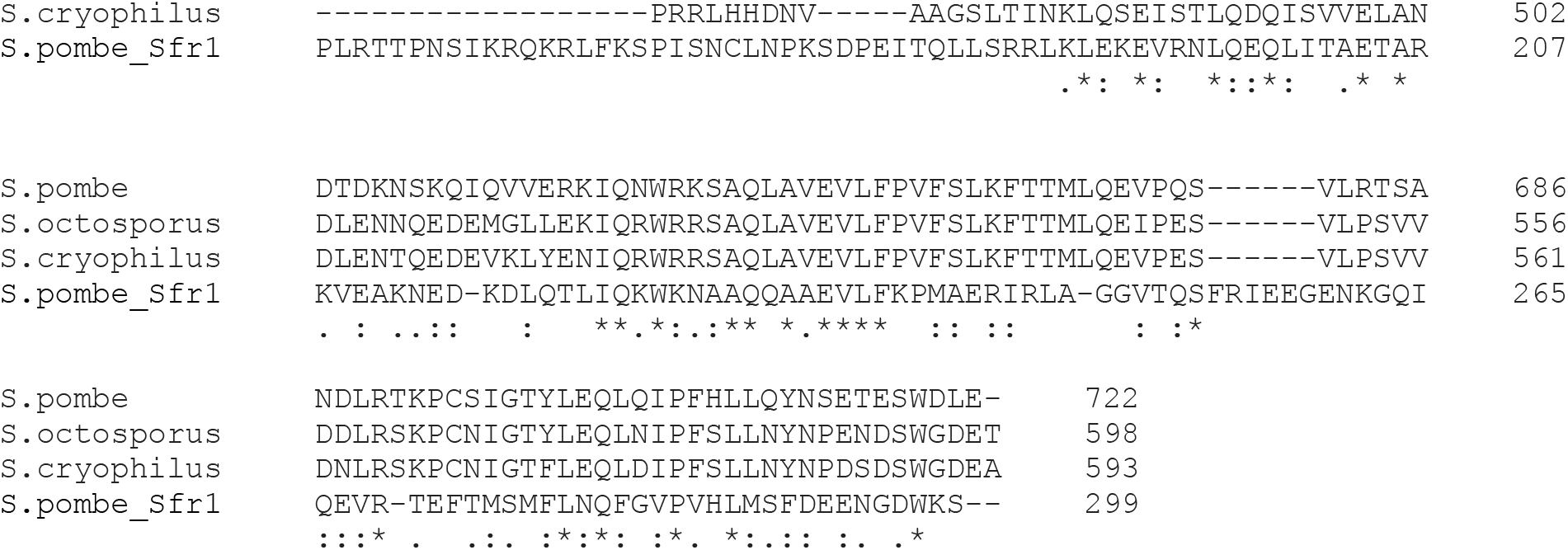
Sequence alignment of Swi2 proteins in fission yeast. Swi2 sequences from *S. pombe, S. octosporus*, and *S. cryophilus* were aligned with Clustal Omega.

**Supplementary Figure S2.**
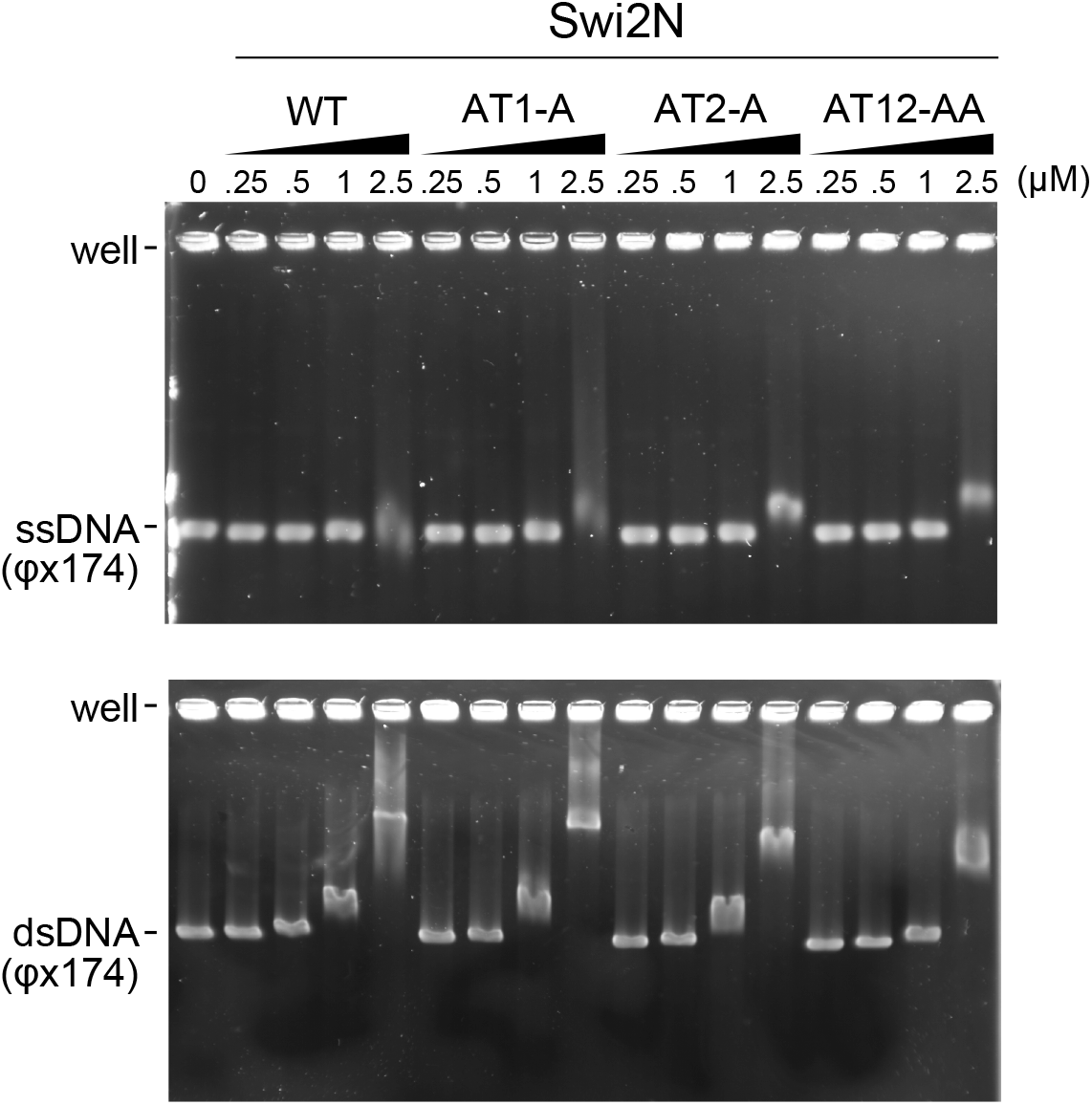
EMSA of Swi2N. EMSA of Swi2N derivatives with a cssDNA (ΦX174 viral DNA) and a circular covalently closed DNA (ΦX174 replicative form DNA).

**Supplementary Figure S3.**
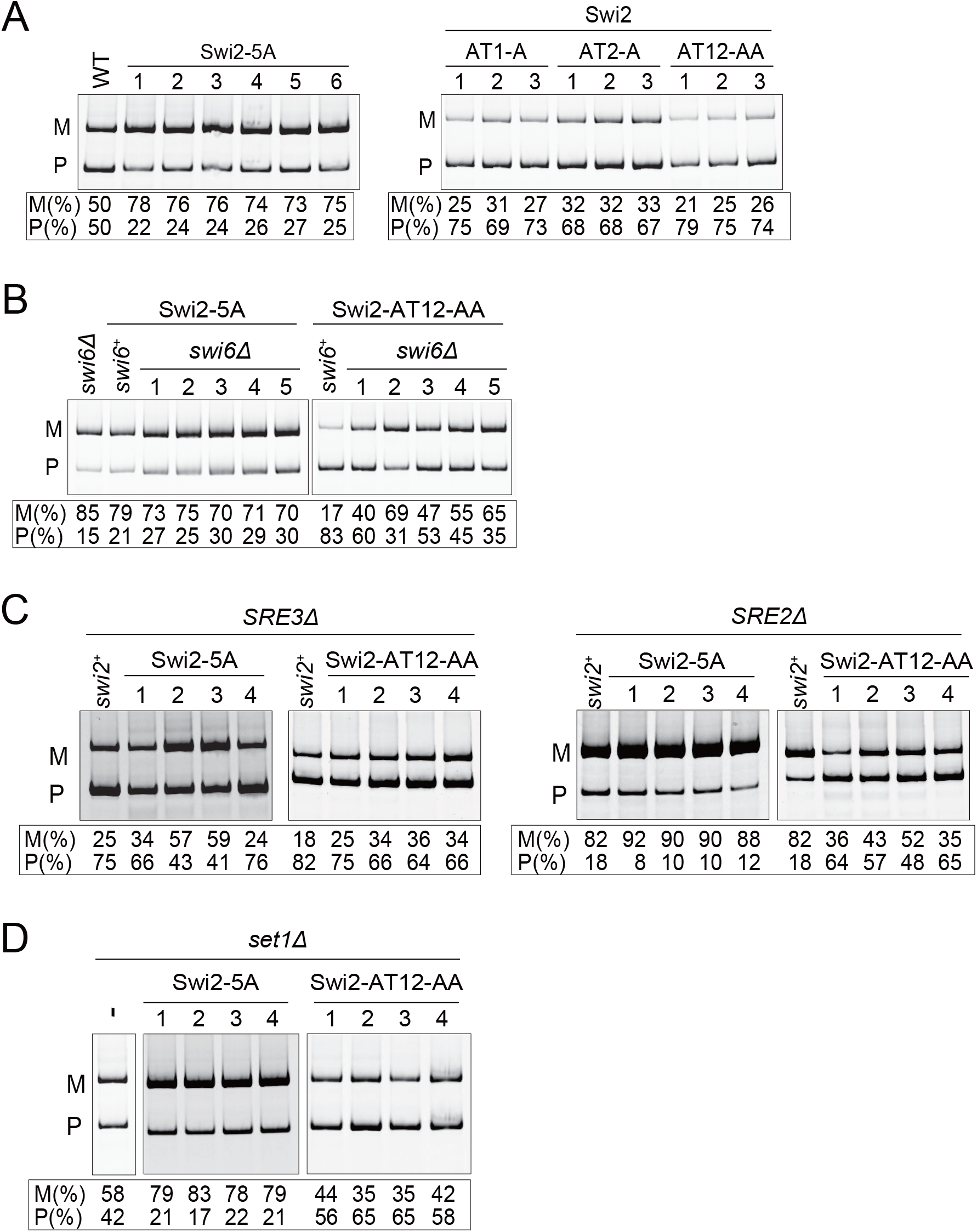
Multiplex PCR analysis of Swi2 mutant cells. (A–D) Gels of multiplex PCR analysis of the Swi2 mutants shown in Figures 4B, 4C, 5B, and 5C. The *mat1-P* and *mat1-M* band intensities were measured, and the percentage of the P or M band intensity was calculated as P/(P+M)×100 or M/(P+M)×100 for each single Swi2 mutant (5A, AT1-A, AT2-A, and AT12-AA) (A), the Swi2-5A *swi6*Δ and Swi2-AT12-AA *swi6*Δ mutant strains (B), the Swi2 mutants combined with deletion of *SRE* elements, *SRE2*Δ or *SRE3*Δ (C), and Swi2-5A *set1*Δ and Swi2-AT12-AA *set1*Δ (D).

## Supplementary Materials For

### The mating-type switching factor Swi2-Swi5 designates a donor through the differential localization modes of Swi2 and stimulation of Rad51-mediated strand invasion in fission yeast *Schizosaccharomyces pombe*

Supplementary Materials including

Supplementary Tables S1 and S2

Supplementary Figures S1-S3

**Supplementary Table 1.** List of Strains used in the study.

| Strain (*S. pombe*) | Mating type region | Other |
| --- | --- | --- |
| EM20 | *h<sup>90</sup>* | *leu1-32 his3-D1 ade6-M375* |
| TM366 | *h<sup>90</sup>* | *swi2-5A leu1-32 his3-D1* |
| TM511 | *h<sup>90</sup>* | *swi2-AT1-AA leu1-32 his3-D1* |
| TM519 | *h<sup>90</sup>* | *swi2-AT2-AA leu1-32 his3-D1* |
| TM527 | *h<sup>90</sup>* | *swi2-AT12-AA leu1-32 his3-D1* |
| TM360 | *h<sup>90</sup>* | *(Blp1)::LEU2 swi6D::ura4 ura4-D18 leu1-32 ade6-216* |
| TM689 | *h<sup>90</sup>* | *swi2-5A-9PK::kamR leu1-32, his3-D1* |
| TM880 | *h<sup>90</sup>* | *swi6D::LEU2 swi2-5A-9PK::kamR leu1-32 his3-D1* |
| M693 | *h<sup>90</sup>* | *swi2-AT12-9PK::kamR leu1-32, his3-D1* |
| TM875 | *h<sup>90</sup>* | *swi6D::LEU2 swi2-AT12-9PK::kamR leu1-32 his3-D1* |
| TM108 | *mat1-PΔ17::LEU2* | *swi2-9PK::kamR* |
| TM107 | *mat1-Msmt-0* | *swi2-9PK::kamR* |
| TM705 | *mat1-PΔ17::LEU2* | *swi2-5A-9PK::kamR ura4-D18 leu1-32* |
| TM711 | *mat1-Msmt-0* | *swi2-5A-9PK::kamR ura4-D18 leu1-32* |
| TM709 | *mat1-PΔ17::LEU2* | *swi2-AT12-AA-9PK::kamR ura4-D18 leu1-32* |
| TM714 | *mat1-Msmt-0* | *swi2-AT12-AA-9PK::kamR ura4-D18 leu1-32* |
| TP75 | *mat3-M SRE3Δ* | *ura4-D18 leu1-32 ade6-M210* |
| TM857 | *mat3-M SRE3Δ* | *swi2-5A-9PK::kamR* |
| TM802 | *mat3-M SRE3Δ* | *swi2-AT12-AA-9PK::kamR* |
| TP8 | *mat2-P SRE2Δ* | *ura4-D18 leu1-32 ade6-M210* |
| TM797 | *mat2-P SRE2Δ* | *swi2-5A-9PK::kamR* |
| TM852 | *mat2-P SRE2Δ* | *swi2-AT12-AA-9PK::kamR* |
| NO47 | *h<sup>90</sup>* | *set1D:: kamR* |
| TM867 | *h<sup>90</sup>* | *set1D::hphR swi2-5A-9PK::kanR leu1-32 his3-D1 ura4-D18* |
| TM868 | *h<sup>90</sup>* | *set1D::hphR swi2-AT12-AA-9PK::kanR leu1-32 his3-D1 ura4-D18* |
| TM19 | *h<sup>90</sup>* | *swi2D::arg3 ura4-D18 leu1-32 his3-D1 arg3-D1* |
| TM100 | *mat1-Msmt-0* |  |
| TM534 | *h<sup>90</sup>* | *swi2-3A leu1-32 his3-D1 ade6-M375* |

| Strain (*S. cerevisiae*) | Mating type | Other |
| --- | --- | --- |
| AH109 | *MATa* | *rp1-901, leu2-3, 112 ura3-52 his3-200 gal4Δ gal80Δ LYS2::GAL1<sub>UAS</sub>-GAL1<sub>TATA</sub>-HIS3 MEL1 GAL2<sub>UAS</sub>-GAL2<sub>TATA</sub>-ADE2 URA3::MEL1<sub>UAS</sub>-MEL1<sub>TATA</sub>-lacZ* |

**Supplementary Table 2.**
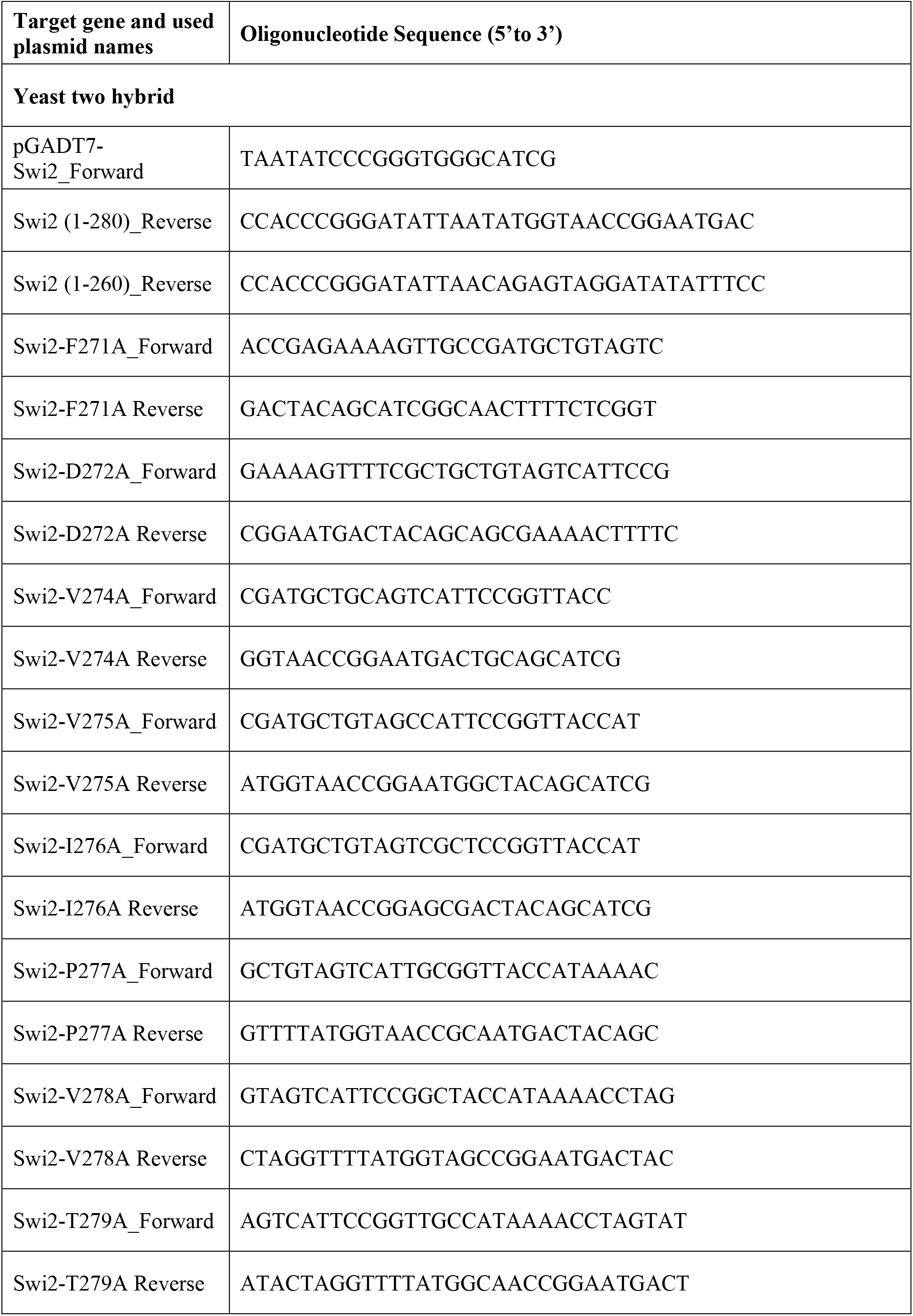
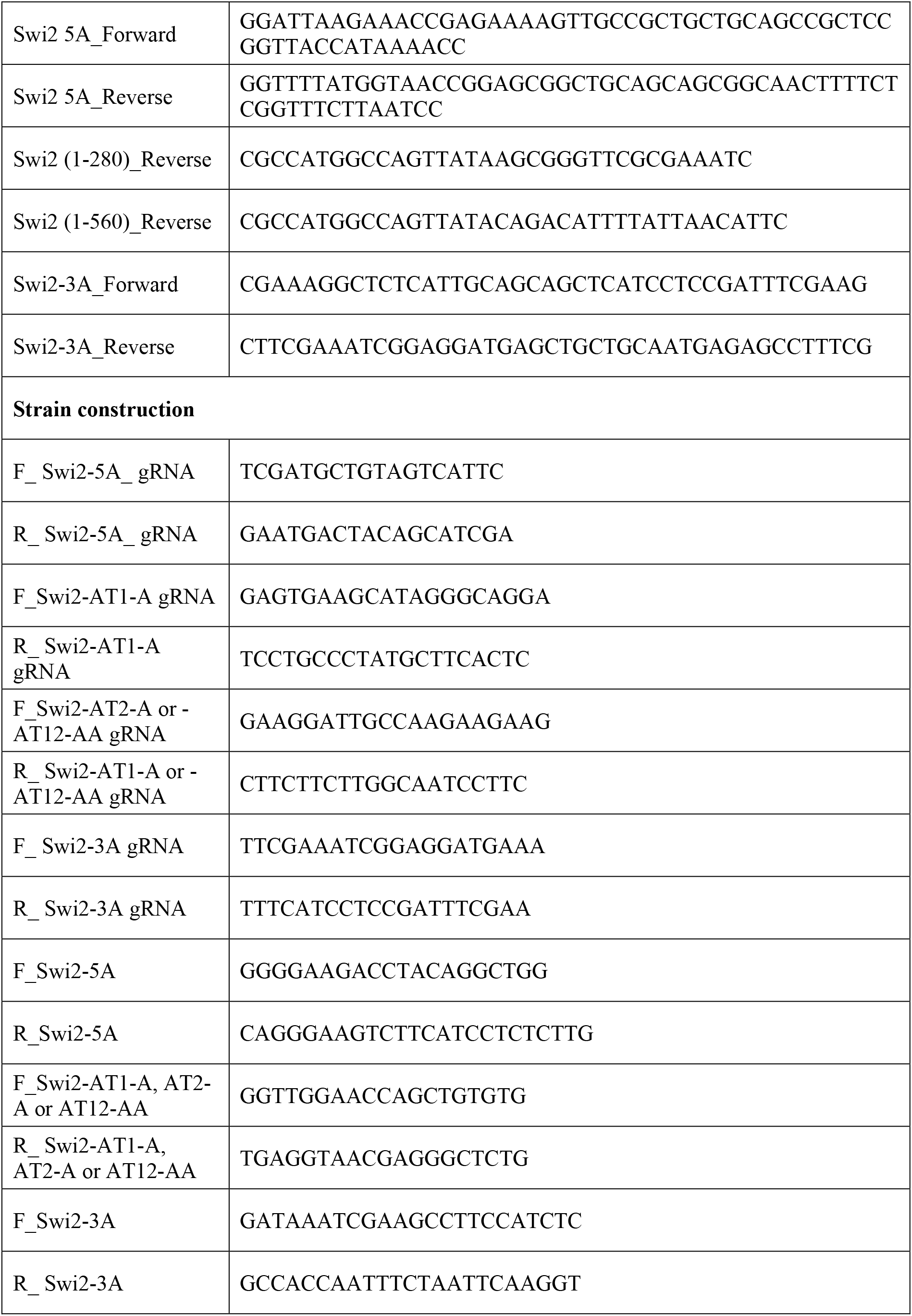
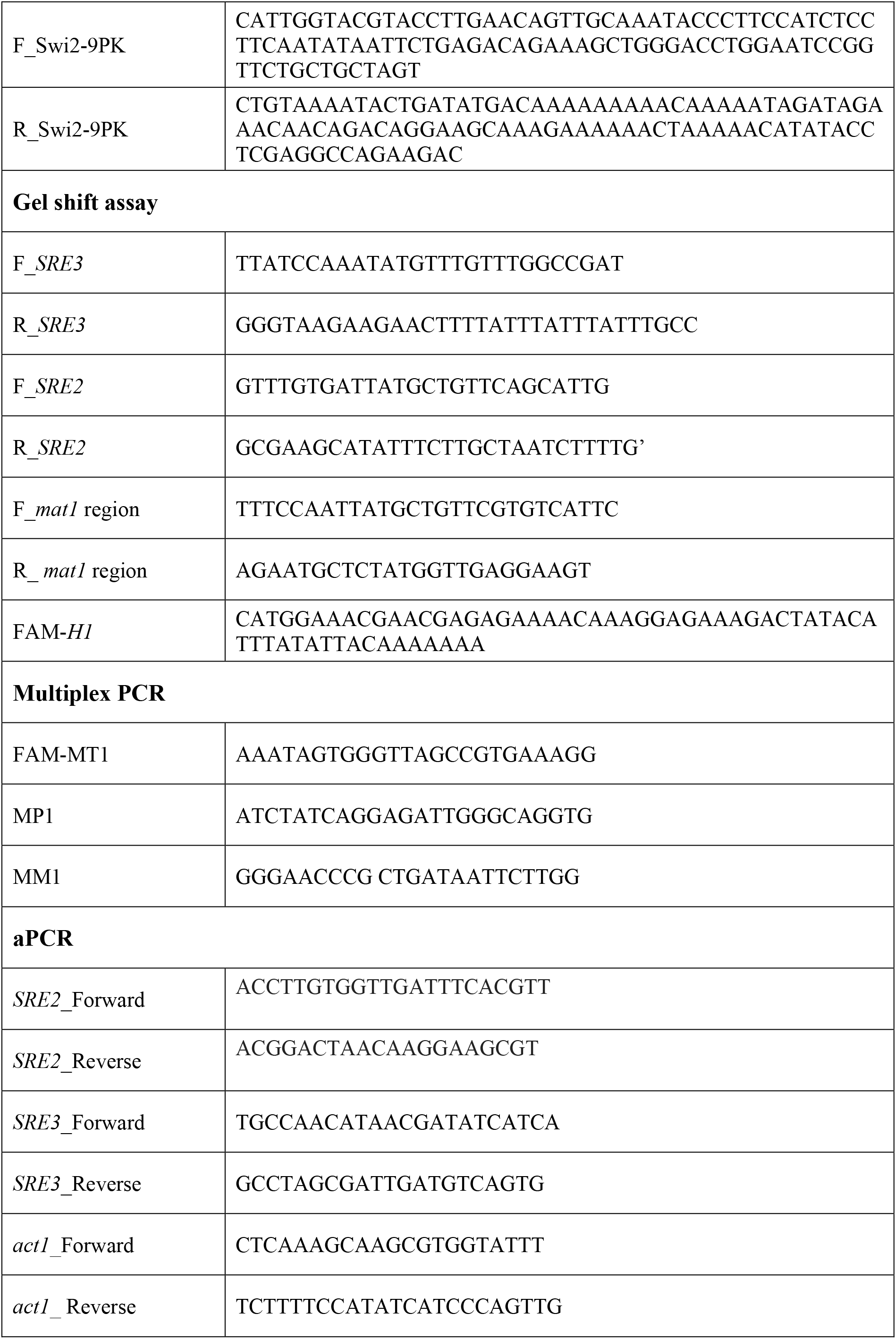
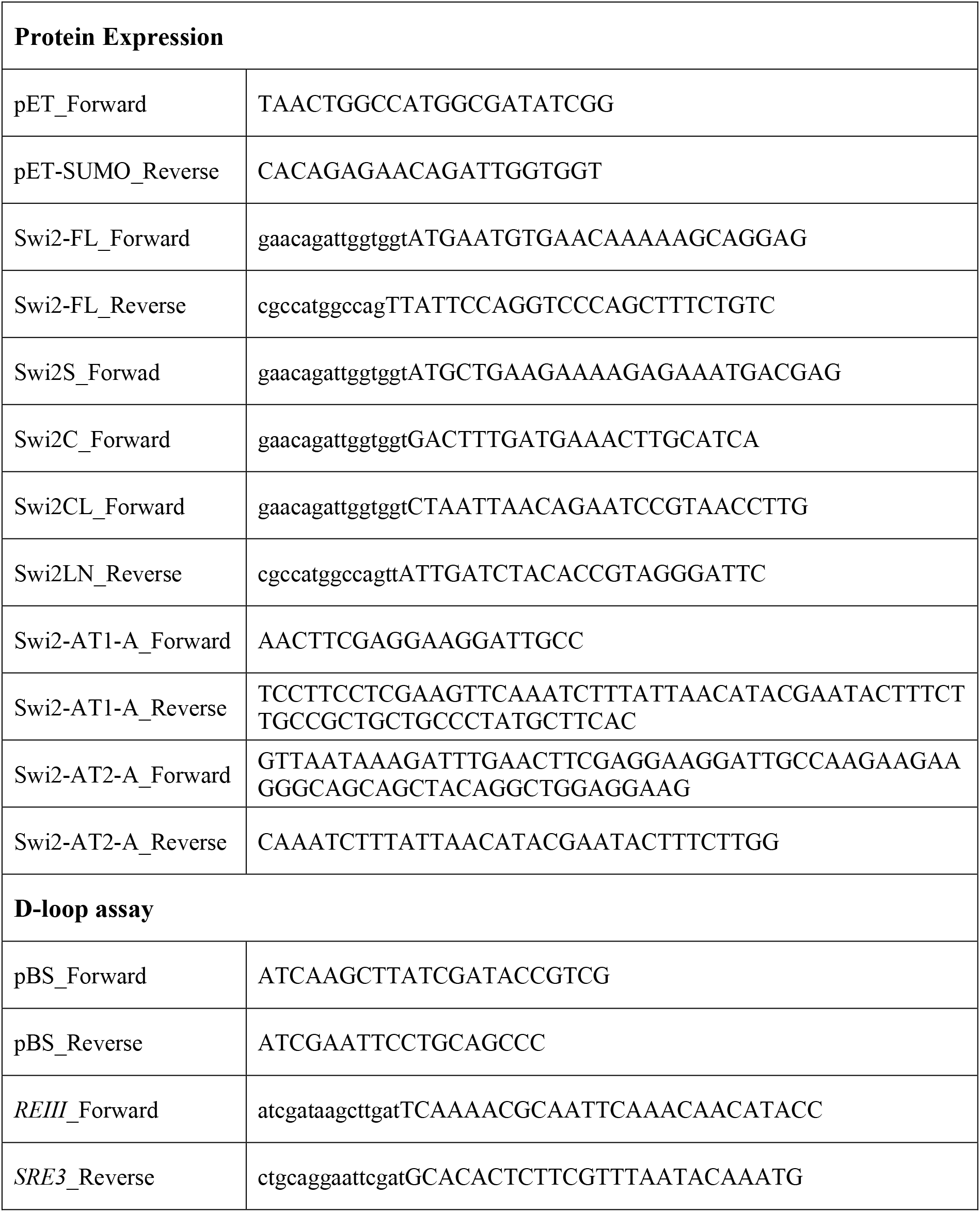
List of Oligonucleotides used in this study.

